# Compromised nuclear envelope integrity drives tumor cell invasion

**DOI:** 10.1101/2020.05.22.110122

**Authors:** Guilherme P.F. Nader, Sonia Agüera-Gonzalez, Fiona Routet, Matthieu Gratia, Mathieu Maurin, Valeria Cancila, Clotilde Cadart, Matteo Gentili, Ayako Yamada, Catalina Lodillinsky, Emilie Lagoutte, Catherine Villard, Jean-Louis Viovy, Claudio Tripodo, Giorgio Scita, Nicolas Manel, Philippe Chavrier, Matthieu Piel

**Affiliations:** Institut Curie and Institut Pierre Gilles de Gennes, PSL Research University, CNRS, UMR 144, Paris, France; Institut Curie, Université PSL, CNRS, UMR 144, Paris, France; Institut Curie, Université PSL, INSERM, U 932, Paris, France; Tumor Immunology Unit, University of Palermo, Corso Tukory 211, 90234, Palermo, Italy; Molecular and Cell Biology Department, University of California, Berkeley, 142 Life Sciences Addition #3200, Berkeley, CA 94720-3200, USA; Broad Institute of MIT and Harvard, 415 Main Street, Cambridge, MA 02142, USA; Institut Curie and Institut Pierre Gilles de Gennes, PSL Research University, CNRS, UMR 168, Paris, France; PASTEUR, Département de chimie, École normale supérieure, Université PSL, Sorbonne Université, CNRS, Paris, France; Research Area, Instituto de Oncología Ángel H. Roffo, Universidad de Buenos Aires, Buenos Aires, Argentina; Consejo Nacional de Investigaciones Científicas y Técnicas (CONICET), Buenos Aires, Argentina; IFOM, the FIRC Institute of Molecular Oncology IFOM. Via Adamello 16 20139 Milano & Department of Oncology and Hemato-oncology, University of Milan. IFOM. Via Adamello 16 20139 Milano, Italy

## Abstract

While mutations leading to a fragile envelope of the cell nucleus are well known to cause diseases such as muscular dystrophies or accelerated aging, the pathophysiological consequences of the recently discovered mechanically induced nuclear envelope ruptures in cells harboring no mutation are less known. Here we show that repeated loss of nuclear envelope integrity in nuclei experiencing mechanical constraints promotes senescence in nontransformed cells, and induces an invasive phenotype including increased collagen degradation in human breast cancer cells, both *in vitro* and in a mouse xenograft model of breast cancer progression. We show that these phenotypic changes are due to the presence of chronic DNA damage and activation of the ATM kinase. In addition, we found that depletion of the cytoplasmic exonuclease TREX1 is sufficient to abolish the DNA damage in mechanically challenged nuclei and to suppress the phenotypes associated with the loss of nuclear envelope integrity. Our results also show that TREX1-dependent DNA damage induced by physical confinement of tumor cells inside the mammary duct drives the progression of *in situ* breast carcinoma to the invasive stage. We propose that DNA damage in mechanically challenged nuclei could affect the pathophysiology of crowded tissues by modulating proliferation and extracellular matrix degradation of normal and transformed cells.

## Introduction

In tissues, extracellular matrix fibers and cell packing limit the available space. Migrating and proliferating cells thus often face situations of crowding and confinement, which can result in strong cell deformations. We and others recently showed that, in the case of migrating immune and cancer cells, strong nuclear deformation can result in transient events of nuclear envelope (NE) rupture with subsequent DNA damage (^1–5^). In this context, efficient repair responses are essential to ensure cell viability, which involve ESCRT proteins restoring NE integrity, as well as BAF1 and the DNA repair pathways (^6^). NE ruptures can be induced by simply confining cells between two plates, independently of their migratory behavior (^4,7,8^). They can also occur spontaneously in fragile mutant nuclei (^9,10^), associated with a large range of degenerative diseases (^11^). Recent studies showed that these events can also occur *in vivo*, in the heart of developing embryos, due to stiffening and beating of the tissue (^12^), as well as in skeletal muscle cells harboring mutations in the lamin A/C gene, due to forces exerted on the nuclei by the microtubule cytoskeleton (^13,14^). Both cases correspond to muscle tissues which are highly solicited mechanically. In these tissues, NE ruptures are associated with senescence and aging phenomena that affect cell proliferation, survival, and ultimately tissue function. NE ruptures are often accompanied by DNA damage of unknown underlying mechanism(s) (^2,3,12,14,15^). Here, we investigated the consequences of nuclear deformation and rupture associated with cell migration in crowded tissues, focusing on mammary duct carcinoma. These tumors, which grow confined inside the mammary ducts, have been recently shown to display large strands of collectively moving cells associated with the transition from *in situ* to the invasive stage of the tumor development (^16^). Additionally, the breast neoplastic invasive switch requires the activity of matrix proteases for breaching out of the duct (^17,18^).

## Results

We first investigated human samples of breast ductal adenocarinoma *in situ* with microinvasion stained by double-marker immunohistochemistry for the small GTPase RAB5A (involved in recycling of cell/cell adhesions ^19^, used as a marker for strands of motile cells^19^) and the DNA damage marker phospho-γH2AX. Nuclei were also stained with Harris’ hematoxylin. We observed that tumor regions positive for RAB5A, often corresponding to micro-invasive foci (see methods and ^20^), displayed a large number of strongly deformed and elongated nuclei (Fig. 1A-D). This morphology is expected in compact strands of collectively migrating cells (^16,21–23^), and corresponds to cells squeezing each other as they move. These deformed nuclei were also more often positive for γH2AX than less deformed ones, and in general displayed more intense γH2AX staining than nuclei in the bulk of the tumor (Fig. 1A-C). This suggests that, at the *in situ* stage of human breast carcinoma, strands of motile cells at the periphery of the tumor exhibit pronounced nuclear deformation, associated with elevated DNA damage.

**Figure 1.**
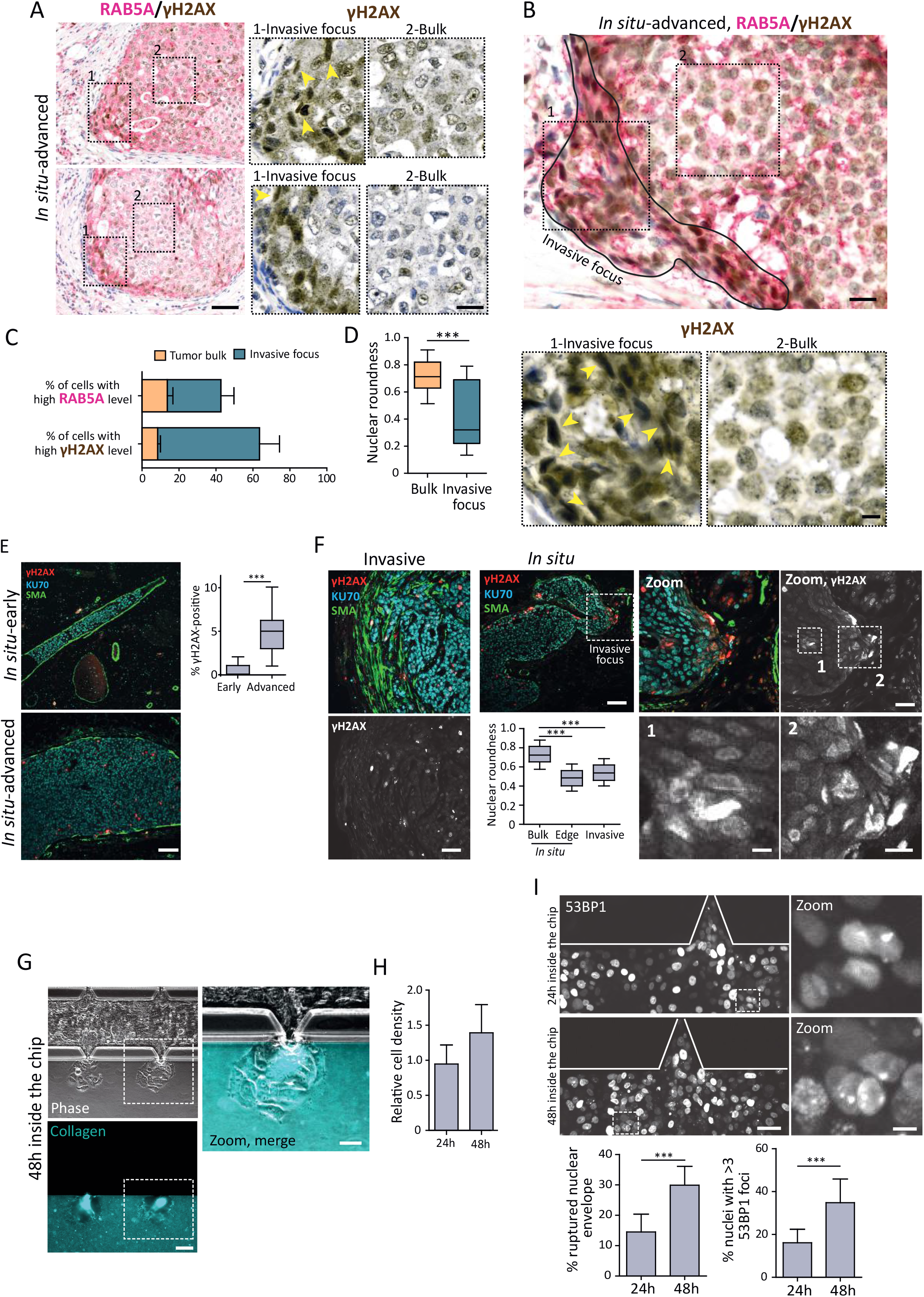
Advanced stages of human breast tumors and mouse xenograft model of breast cancer progression display strong nuclear deformation and DNA damage. (A) Immunohistochemistry of *in situ* advanced human primary breast tumors. RAB5A (red), γH2AX (brown). Yellow arrowheads point to highly deformed nuclei. Bar, 80 μm. (B) Magnified region of *in situ* advanced primary breast tumor with a micro-invasive focus. Yellow arrowheads point to highly deformed nuclei. Bar, 25 μm. (C) Quantification of RAB5A and γH2AX intensity levels in the tumor bulk and invasive foci. Data are the mean ± SD. (D) Box and whisker plot showing the median value and 10-90 percentiles of the nuclear roundness index measured in the tumor bulk and invasive foci. (A-D) Images and quantifications are representative of 7 independent tumors samples where 21 and 27 nuclei were analyzed per sample for tumor bulk and invasive foci, respectively. (E, F) Immunofluorescence analysis of alpha-smooth muscle actin, SMA (green), human-specific KU70 (cyan) and γH2AX (red) in 7 weeks old mouse xenografts at different stages (*in situ*-early, *in situ*-advanced, *in situ* with invasive focus and Invasive) generated by intraductal injection of DCIS cells. (E) The *in situ*-advanced stage is characterized by a discontinuous and broken myoepithelial layer, as opposed to the *in situ*-early stage, which exhibits a continuous myoepithelial layer. Graph, box and whisker plot showing the median value and 10-90 percentiles of the percentage of γH2AX-positive nuclei measured at *in situ*-early and -advanced stages. Data represents 3 independent experiments where 500 nuclei were scored per tumor stage, per experiment. (F) Invasive stage: SMA-positive and elongated cells represent cancer associated fibroblasts. Graph, box and whisker plot showing the median value and 10-90 percentiles of the nuclear roundness index measured at invasive and *in situ* (further divided into edge,up to 2 cell layers below the myoepithelial layer, and bulk) tumor stages. Data represents 3 independent experiments where 7 *in situ* and 13 invasive tumor samples were analyzed (for a total of 140 nuclei analyzed for *in situ*-bulk, 139 nuclei analyzed for *in situ*-edge and 264 nuclei analyzed for invasive tumor samples). (E, F) Bar, 100 μm, (F) Zoom bar, 50 μm; inset bars, 15 μm. (G) Duct-on-chip assay. Images were acquired 48 h post-cell injection. Alexa fluor 647-labelled type I collagen was mixed with unlabeled type I collagen (blue). (H) Graph, cell density was measured by counting the number of cells in 5 random fields (20x magnification) at the indicated time points. Data represents the mean ± SD of 3 independent experiments. Bar, 50 μm; zoom bar, 25 μm. (I) Duct-on-chip assay. DCIS cells stably expressing 53BP1-mCherry (gray levels on the shown image) and cGAS-EGFP were injected in the chip device. Images were acquired at the indicated time points to measure DNA damage levels (number of 53BP1 foci) and NE rupture events. Data represents the mean ± SD of 3 independent experiments. Bar, 40 μm; zoom bar, 10 μm. *P* values were calculated by unpaired Student’s *t*-test, ***P < 0.0001.

We then investigated a xenograft mouse model based on the intraductal (intra-nipple) injection of transformed human breast MCF10DCIS.com (DCIS) cells, which are derived from the nontransformed MCF10A cell line, and that recapitulates the transition from *in situ* to invasive stages of breast cancer progression (^17,24^). Tumors at various stages were analyzed. *In situ* tumors (further divided in early and advanced stages based on the continuity of the myoepithelial layer) are characterized by a myoepithelial layer (stained with smooth muscle actin) surrounding human xenograft cells that were identified by the human-specific KU70 antibody. In order to verify whether human tumor cells similarly experience DNA damage in the tumor xenografts, tumor sections were stained with γH2AX (Fig. 1E-F, Fig. S1A). Similar to human tumor specimens, xenograft tumors displayed numerous deformed nuclei, mostly found at the periphery of the tumor mass, close to the myoepithelium, especially in ducts inflated with large tumors (Fig. 1E-F), and enriched in regions showing micro-invasive foci. Strikingly, these regions were also enriched in γH2AX-positive nuclei (Fig. 1F). Since γH2AX is also known to stain mitotic cells, a detailed investigation of the DAPI staining revealed that most of the γH2AX-positive cells did not correspond to mitotic figures with a highly condensed DNA content, which are distinct from DNA damage-positive cells (Fig S1A). Together, these data show that mammary *in situ* carcinomatous lesions, both in human and in xenografted mice often display strongly deformed nuclei with elevated levels of DNA damage, possibly due to NE ruptures, as suggested by our previous work on migrating cells (^2,3^).

Because NE ruptures of deformed nuclei are difficult to investigate in tumor samples, we developed a ‘duct-on-chip’ microfluidic assay (inspired by ^25^). The assay consists of three large 50 μm-high channels: a lateral channel filled with culture medium, a central one with DCIS cells, and the third channel with a dense fluorescently-labeled type I collagen matrix. The channels are connected by regularly spaced small V-shape gates of 10 μm-wide smaller section through which cells can migrate towards the collagen chamber (Fig. 1G and Fig. S1B). Cells were able to invade efficiently the collagen chamber (Fig. 1G) and visualization of the collagen network suggested that cells were able to remodel and possibly degrade the collagen fibers over time (Fig. 1G). Indeed, using the pan inhibitor of matrix metalloproteinases (MMPs), GM6001, we confirmed that collagen degradation by DCIS cells was required for invasion into the collagen chamber (Fig.S1C, D), similar to what we have observed *in vivo* (^17^). This confirms that the ‘duct on a chip’ model, although simplified, can be used to mimic the intraductal space and thus can recapitulate key features of the *in situ*-to-invasive transition of ductal breast carcinoma (^25^).

DCIS cells expressing catalytically inactive cGAS-mCherry (hereafter referred to as icGAS) to detect NE ruptures (^3,8^), and 53BP1-GFP to detect DNA double strand break foci were grown in the duct-on-chip device. When comparing cells at 48 *vs* 24 h after injection in the device, the average cell density increased by 50% (Fig 1H) and the average fraction of cells displaying NE ruptures and DNA damage doubled (Fig. 1I), suggesting that increased cell density could be responsible for crowding-induced nuclear deformation and rupture. Strikingly, DCIS cells remained highly motile even at high density (Movie S1), confirming previous observations (^16,19^). This suggests that, similarly to what was observed for single migrating cells in a dense tissue (^2,3^), the motility of DCIS cells in a confined and crowded multicellular context can lead to nuclear deformation and rupture and is accompanied by increased DNA damage.

To examine more precisely the link between nuclear deformation, NE rupture and DNA damage, we used a 2D confiner device that allows precise deformation of cells between two parallel plates (^26^). In addition to DCIS cells, we analyzed the non-malignant cell lines RPE1 (normal human hTERT immortalized cells, in which we have previously characterized NE ruptures and DNA damage in migration assays in ^3^) and MCF10A (normal human breast epithelial cells, the parental of the DCIS cell line). Using cells expressing icGAS-mCherry and 53BP1-GFP (Figure 2A), we found a sharp increase in the fraction of cells showing ruptured nuclei below 3 μm confinement (Fig. S2A, B), confirming our previous observations on another panel of cell types (^3^). Importantly, only cells with ruptured NE displayed increased number of 53BP1 foci (Fig. 2A, B and Fig. S2C, D). Cells exhibiting NE ruptures were also positive for other DNA damage markers (Fig S2E, F). Importantly, the elevated number of 53BP1 foci remained high throughout the confinement period (Fig. 2C and S2D). This was associated with a majority of cells showing repeated cycles of NE blebbing, rupture and repair for the entire duration of the confinement (Movie S2). These findings indicate that long confinement periods could lead to an extended exposure of nuclear chromatin to DNA-damaging factors and thus elevated levels of DNA damage. In addition, these experiments revealed that in RPE1, MCF10A and DCIS cells, there was a strong correlation between NE rupture events and a sustained elevated number of DNA damage foci for the entire duration of the confinement. Altogether, these observations suggest that cells undergoing strong nuclear deformation within tissues might experience chronic DNA damage for long periods of time.

Previous work suggested that NE rupture could lead to increased levels of DNA double strand breaks due to a defect in the DNA repair process as a result of the loss of DNA repair proteins out the nucleus upon NE rupture (^1^). An alternative hypothesis is that some cytoplasmic factor(s) (proteins or other molecules) could get inside and access the nuclear DNA, causing DNA damage (^27–29^). We quantified the number of DNA damage foci appearing per unit of time and their lifetime in RPE1 cells expressing 53BP1-GFP, comparing cells treated with the topoisomerase inhibitor etoposide to cells confined as above (in cells experiencing NE rupture or not). We found that the appearance of new foci per unit of time was higher in cells with NE rupture, reaching almost the same rate as in the etoposide-induced case (Fig 2D). Moreover, the number of new foci often increased only after a rupture event (Fig S2G, H) suggesting a causal relationship between NE rupture and new DNA damage. On the contrary, lifetime of the foci was only slightly increased in confined cells compared to etoposide-treated cells and showed no difference between cells with or without NE ruptures. This suggests that DNA repair was slower in confined cells but not significantly affected by NE rupture (Fig. 2E, the duration of NE repair was estimated to be only a few minutes, much shorter than the DNA repair process^3^). Overall, these experiments suggest that DNA damage associated with NE ruptures is mostly due to the generation of new DNA damage following NE rupture events rather than delayed repair and that the mechanism producing this damage remains active in the nucleus for a long time following NE repair.

**Figure 2.**
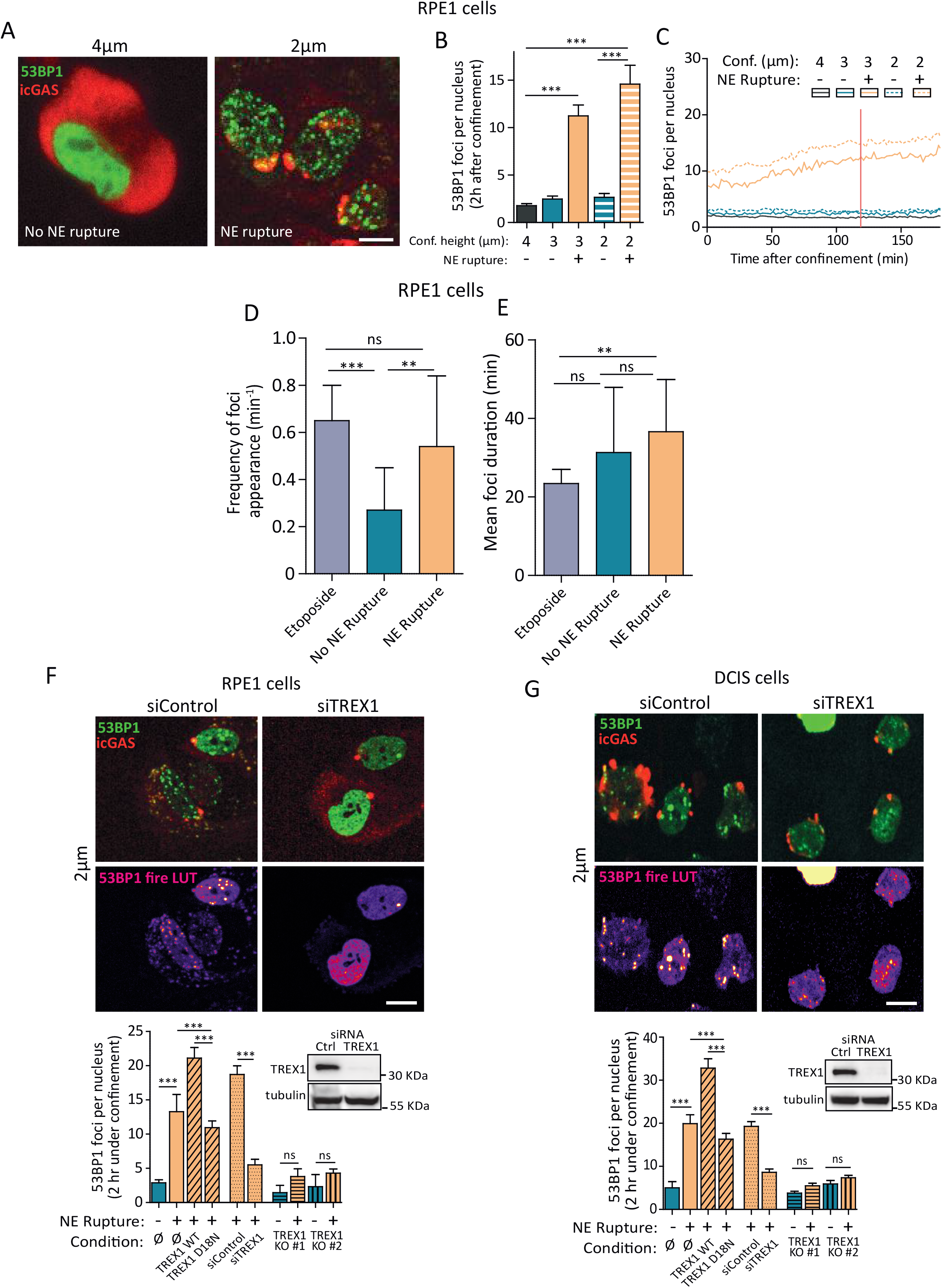
Strong confinement induces nuclear envelope rupture and TREX1-dependent DNA damage. (A) RPE1 cells stably expressing 53BP1-EGFP and catalytically inactive cGAS-mCherry were confined at the indicated heights and images were acquired while cells were under confinement (after 2 h). Bar, 5 μm. (B, C) Quantification of DNA damage levels during confinement as assessed by the number of 53BP1 foci in cells displaying or not NE rupture. Data represents the mean ± SEM of 3 independent experiments where 20 cells per experiment and per condition were analyzed. Red vertical line indicates a single time-point, which is represented in “B”. (D, E) Frequency of foci appearance and foci life-time (foci duration) for RPE1 cells confined at 2 μm (with or without NE rupture) or treated with etoposide (25 μM). Data represents the mean ± SD of 3 independent experiments where 20 cells per condition were analyzed. (F) A variety of TREX1-deficient or -proficient RPE1 cells (all stably expressing 53BP1-EGFP and catalytically inactive cGAS-mCherry) were confined at 2 μm height and DNA damage levels were assessed after 2h under confinement by the number of 53BP1 foci in cells displaying or not NE rupture. (G) Same as in “F” but with DCIS cells. (F, G) Data represents the mean ± SEM of 3 independent experiments where 20 cells were analyzed per condition per experiment. Western blots show TREX1 depletion in RPE1 and DCIS cells 48h post-knockdown; tubulin is the loading control. Bar, 10 μm. *P* values were calculated by unpaired Student’s *t*-test, ***P < 0.0001; **P < 0.005; ns = not significant.

NE ruptures at chromatin bridges that persisted after mitotic exit due to lagging chromosomes have been shown to cause DNA damage (^28,29^). Importantly, DNA damage was reduced upon depletion of TREX1, a cytoplasmic ER membrane-associated exonuclease (^28^). Thus, we tested the TREX1-dependency of DNA damage events occurring in cells under confinement. We used the same collection of normal and transformed cell lines (RPE1, MCF10A, DCIS) and TREX1 was either transiently silenced by siRNA-mediated knockdown, or permanently knocked out by CRISPR technology (RPE1 and DCIS-Fig. S2I, J). Strikingly, we found that NE ruptures were not accompanied by an increase in 53BP1 foci in cells transiently depleted for TREX1 by siRNA and in knockout cells (Fig 2F, G and Fig. S2C, D). TREX1 depletion did not affect the generation of 53BP1 foci upon etoposide treatment (Fig S2K) and TREX1 knockout clones displayed NE rupture rates comparable to the parental cell line (Fig. S2L). Overexpression of wild-type TREX1 increased the number of 53BP1 foci in cells with NE ruptures, while overexpression of a catalytically-deficient mutant (TREX1-D18N) (^30^) decreased foci generation (Fig 2F, G). We confirmed that, in non-confined cells, TREX1-GFP is mostly bound to endomembranes and excluded from the nucleoplasm. Upon confinement, TREX1-GFP foci were also found inside the nucleus (Fig S2M, Movie S3). Nevertheless, we could not clearly observe a translocation of TREX1-GFP through NE rupture sites. Overall, these experiments establish TREX1 depletion as an efficient way to prevent elevated damage upon NE rupture in confined cells, although the mechanism of entry and action of TREX1 inside the nucleus remains largely unknown.

In non-transformed cells, chronic exposure to DNA damage-inducing factors is associated with cellular senescence. In agreement, RPE1 and MCF10A cells showed a very clear induction of senescence upon treatment with DNA damaging agents (etoposide and doxorubicin, hereafter doxo) as verified by senescence-associated-β-galactosidase (β-gal)-positive staining. However, senescence was less pronounced in DCIS cells (Fig S3A). Our observations pointed to recurrent DNA damage as long as cells remained confined under heights that induced robust NE rupture (Fig. 2C and S2D). Therefore, we asked whether strong and long-lasting confinement could induce cellular senescence. First, we investigated the cell division cycle of confined cells. RPE1 cells were confined for either a short period (3 h) or a long period (12 h), at heights inducing (2 μm) or not (4 μm) elevated rates of NE ruptures. We observed that during the confinement period, even for long confinement periods, most cells had not undergone mitosis, nor shown any sign of death (Fig S3B, C). Following the confinement periods, we harvested the cells and replated them in order to track single cells that had ruptured their NE or not (based on icGAS perinuclear staining/distribution, Fig S3D). After a short period of confinement, de-confined cells resumed proliferation with an almost normal cell cycle duration (Fig. 3A, mean cell cycle duration for non-confined cells: 14.66 h +/− 2.31), regardless of their NE status. Similarly, cells confined for a long period without NE rupture exhibited only a slight increase in cell cycle duration (Fig 3A). In contrast, cells that displayed NE rupture and that had been confined for a long period, while showing no sign of death, did not resume proliferation (Fig. 3A right panel) and for the few dividing cells, showed a much longer cell cycle duration (Fig 3A). These results suggest that NE rupture and sustained confinement induced DNA damage delay cell cycle progression, indicative of the activation of a cellular senescence program.

**Figure 3.**
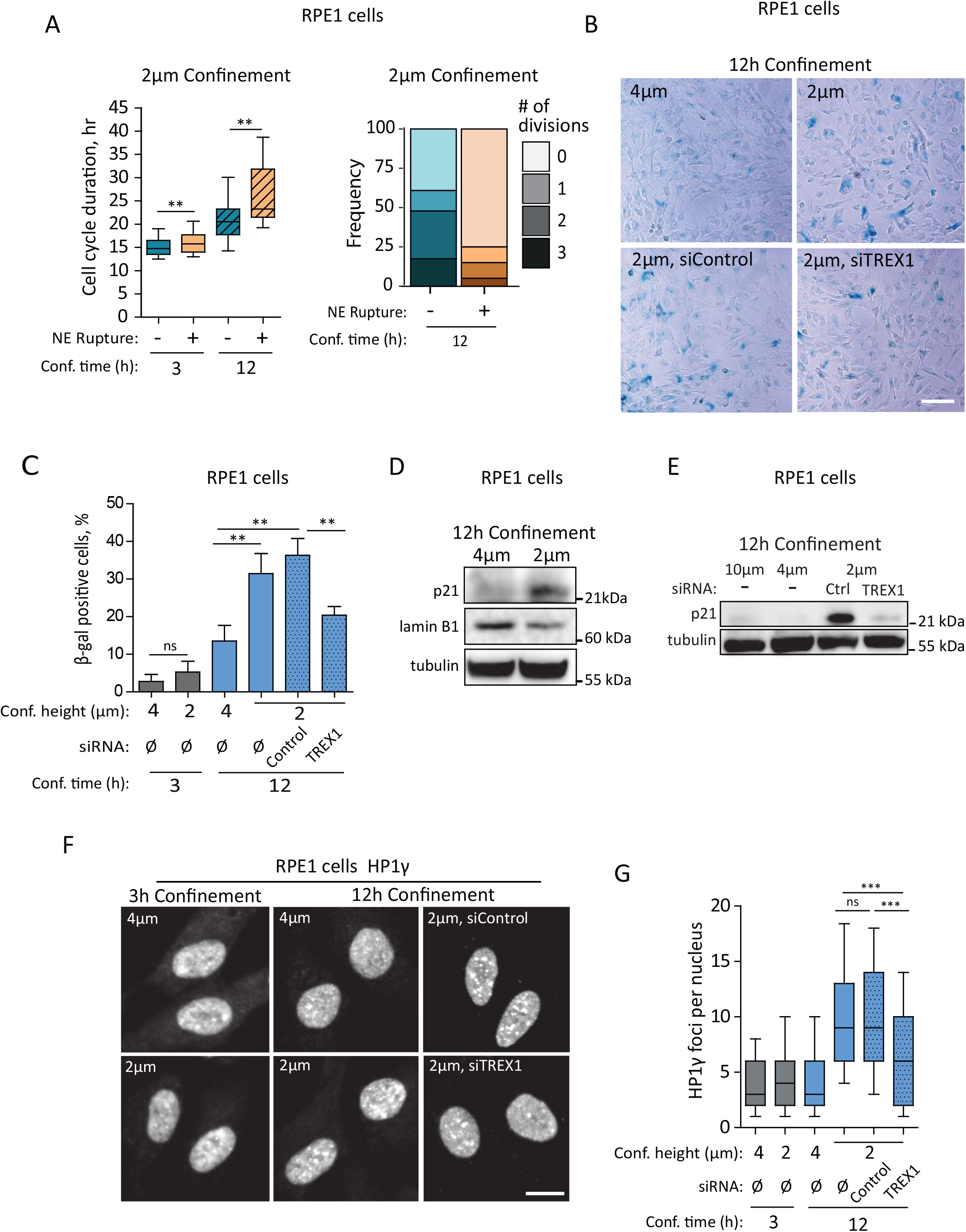
Long-term, strong confinement causes cell senescence in a TREX1-dependent manner. (A) RPE1 cells stably expressing catalytically inactive cGAS-EGFP were confined at 2 μm for either 3 or 12 h. Following these confinement periods, the confinement lid was removed and cells were harvested and replated for imaging for 72 h for cell cycle duration measurement of cells displaying or not NE rupture. Graph, left: box and whisker plot showing the median value and 10-90 percentiles of the cell cycle duration. Graph, right: frequency distribution of the number of cell divisions for cells confined for 12 h at 2 μm. Data for 3 h of confinement represents 3 independent experiments where a total of 150 cells with nonruptured and 200 cells with ruptured nuclei were analyzed. Data for 12 h of confinement represents 3 independent experiments where a total of 23 cells with non-ruptured and 40 cells with ruptured nuclei were analyzed. (B-G) RPE1 cells were transiently depleted for TREX1 using siRNA and confined at the indicated heights for the indicated periods. Following the confinement periods cells were harvested, replated and cultured for 72 h before fixation for β-gal staining (B, C) and immunostaining of heterochromatin foci (HP1γ) (F, G), or lysis for western blot analysis of lamin B1 and p21. GAPDH is the loading control (D, E). (C) Graph, data represents the mean ± SD of 3 independent experiments where 200 cells were scored per condition per experiment; western blot images are representative of 2 independent experiments. (G) Graph: box and whisker plot showing the median value and 10-90 percentiles of the number of HP1γ foci per nucleus. Data represents 3 independent experiments where 150 cells were scored per condition per experiment. Bars, 80 μm (B) and 10 μm (F). *P* values were calculated by unpaired Student’s *t*-test, ***P < 0.0001; **P < 0.005; ns = not significant.

Therefore, we further sought to characterize the senescence phenotype associated with strong confinement. Cells were confined for short and long periods of time, at low (4 μm) or strong (2 μm) confinement, then harvested and replated for 96 h before testing a panel of senescence markers (β-gal-positive staining, heterochromatin foci-HP1γ staining, western blot for lamin B1 and p21 ^31^). All the markers consistently showed that, while short term, or weak (4 μm) confinement did not induce senescence, long term, strong confinement, induced a robust senescence phenotype, in both RPE1 and MCF10A, but not in DCIS cells (Fig 3B-G and S3E-G). Importantly, this senescence phenotype was TREX1-dependent, as TREX1-depleted RPE1 cells showed no senescence even for strong and prolonged confinement (Fig. 3B-G). This shows that, unexpectedly, the mechanism underlying the induction of cell senescence by strong and prolonged confinement is specifically due to TREX1-dependent DNA damage in cells with NE ruptures. Importantly, the senescence phenotype did not require the activation of the canonical cGAS pathway (^31^), since RPE1 cells do not express cGAS (Fig S3H, ^32^). Together, these results show that TREX1-dependent chronic DNA damage induced upon strong confinement triggers hallmarks of senescence in normal RPE1 and MCF10A cells, but not in transformed DCIS cells.

DCIS cells did not show any senescence phenotype associated with strong and prolonged confinement, even if these conditions induced high level of sustained DNA damage. To recapitulate and investigate the potential consequences of DNA damage associated with nuclear deformation and NE ruptures during the development of tumors produced by DCIS cells, we used the duct-on-chip assay described above. When growing under intraductal-like confinement inside the duct-on-chip model, DCIS cells exhibited increased DNA damage after 48 h inside the device (Fig 1I) and TREX1-deficient DCIS cells displayed reduced DNA damage upon strong nuclear deformation under confinement (Fig 2G). Therefore, we asked whether TREX1 was also responsible for the DNA damage observed in cells growing inside the duct-on-chip system. Both control- and TREX1-depleted cells reached high density within 48 h after injection in the device, showing that proliferation was not affected by TREX1 depletion. They also displayed comparable levels of NE ruptures (Fig. 4A). After 24 h in the device, there was no appreciable difference in 53BP1 foci numbers between control- and TREX1-depleted cells (Fig. 4A). In contrast, while this number strongly increased at 48 h for control cells, it remained constant in TREX1-depleted cells (Fig. 4A). These results show that the increased 53BP1 foci levels observed at high density in the duct-on-chip system is a consequence of TREX1-dependent DNA damage following nuclear deformations and ruptures.

**Figure 4.**
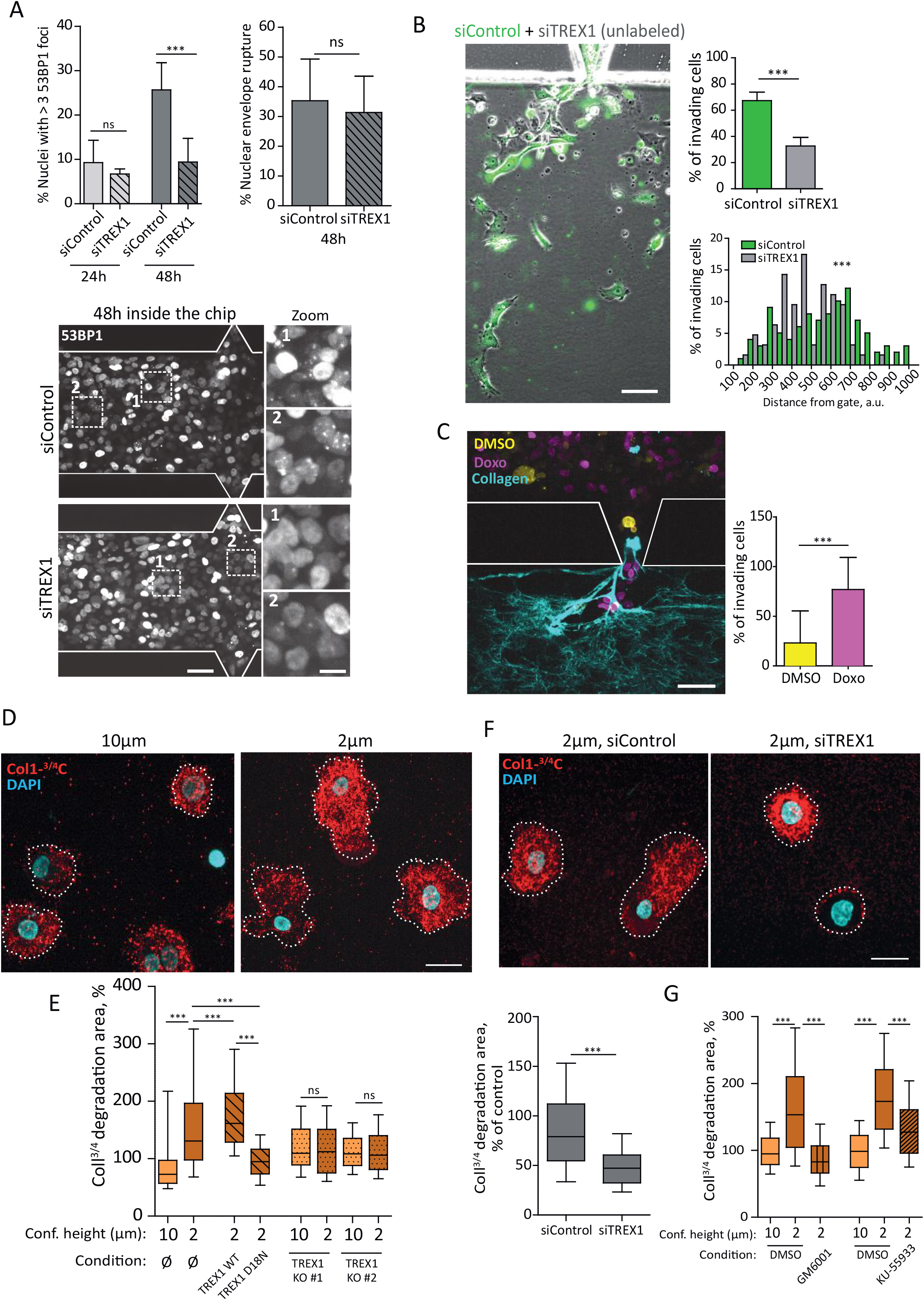
TREX1-dependent DNA damage upon strong confinement leads to collagen degradation and invasion. (A) Duct-on-chip assay. DCIS cells stably expressing 53BP1-mCherry (gray levels) and catalytically inactive cGAS-EGFP were transiently depleted for TREX1 using siRNA and 48 h later injected in the chip device. Images were acquired at the indicated time points to measure DNA damage levels (number of 53BP1 foci) and NE rupture events. Graphs: data represents the mean ± SD of 3 independent experiments where 5 random fields (150 cells scored per field) were analyzed per condition per experiment. Bar, 50 μm; zoom bar, 10 μm. (B) Duct-on-chip assay. DCIS cells were transiently depleted for TREX1 using siRNA and 48 h later injected in the same chip device with siControl cells (prelabled with a cell tracker-green). Graphs: data represents the mean ± SD of 4 independent experiments where cells invading from 7 random gates were scored per experiment. Bar, 50 μm. (C) Ducton-chip assay. DCIS cells stably expressing 53BP1-EGFP were pre-treated for 12 h with doxorubicin (doxo, 20 nM) and mixed with DCIS cells stably expressing 53BP1-mCherry that had been pre-treated for 12 h with DMSO (vehicle). The mixed cell population was then injected in the chip device and pictures were taken 24 h later. Graph: data represents the mean ± SD of 2 independent experiments where cells invading from 10 random gates were scored per experiment. Bar, 40 μm. (D) DCIS cells were confined for 2 h at the indicated heights. Following the confinement period, cells were harvested and embedded in 3D type I collagen for 12 h to assess their collagen degradation activity by immunostaining with an antibody that specifically recognizes the collagenase-cleaved ¾ fragment of collagen I (red); DAPI (cyan). (E) A variety of TREX1-deficient or -proficient DCIS cells were confined at the indicated heights and collagen degradation area was measured. Graph: box and whisker plot showing the median value and 10-90 percentiles of the collagen degradation area. Data represents 3 independent experiments where 50 cells per condition per experiment were analyzed. (F) It was proceeded exactly as described in “D” except that it was performed with TREX1-depleted DCIS cells. Graph: box and whisker plot showing the median value and 10-90 percentiles of the collagen degradation area. Data represents 3 independent experiments where 60 cells per condition per experiment were analyzed. (G) It was proceeded exactly as described in “D” except that cells were treated with the ATM inhibitor (KU-55933, 10 μM) or MMP pan inhibitor (GM6001, 40 μM) while under confinement and also when embedded in 3D type I collagen. Graph: box and whisker plot showing the median value and 10-90 percentiles of the collagen degradation area. Data represents 3 independent experiments where 60 cells per condition per experiment were analyzed. *P* values were calculated by unpaired Student’s *t*-test, ***P < 0.0001; ns = not significant.

TREX1-depleted cells can thus grow at high density in the duct-on-chip device without showing an increase in DNA damage. This prompted us to investigate whether such reduced DNA damage would impinge on the ability of these cells to invade the collagen in the duct-on-chip assay. To compare the behavior of control- and TREX1-depleted cells, we pre-labeled control DCIS cells with a cell tracker (green) and mixed them with an equal number of non-labelled TREX1-depleted cells prior to injection into the chip. We observed that control cells could invade the collagen chamber earlier and migrated farther than TREX1-depleted cells (Fig. 4B). This suggests that the increased invasive behavior of control cells compared to TREX1-depleted cells might be due to the fact that they experience more DNA damage. To test this hypothesis, differently-labelled control and doxo-treated cells (12 h of pre-treatment with doxo, in order to induce long-term, chronic DNA damage) were mixed and introduced in the same chip. We observed that, 24 h after loading the cells into the device (to ensure cell density was still low) doxo-treated cells displayed increased invasion capacity into the collagen chamber as compared to DMSO-treated cells (Fig 4C). These results indicate that sustained DNA damage can be a driver of the invasive behavior of DCIS cells in a 3D matrix environment. This in turn suggests an intriguing hypothesis that confined cell growth due to cell crowding, might favor cell invasion by causing nuclei deformation, which leads to NE ruptures and TREX1-dependent DNA damage.

A major determinant of the ability of cells to escape the central confining chamber and invade the collagen compatment in the duct-on-chip assay is the capacity of the cells to degrade collagen (Fig. 1G and S1C, D). Collagen degradation is also a requirement in the DCIS intraductal xenograft tumor model for the transition from *in situ*-to-invasive carcinoma (^17,18^). We thus assessed the collagen degradation activity of DCIS cells embedded into a 3D type I collagen matrix for 12 h and stained with an antibody that specifically recognizes the collagenase-cleaved ¾ fragment of collagen I (^17,33,34^). Importantly, cells were treated with drugs, or confined for several hours, and then were embedded in the 3D collagen gel in the absence of any further DNA damaging condition. This assay was thus testing the capacity of cells, that have been subjected to sustained DNA damage or not, to maintain a collagenolytic activity for the following hours. We first treated DCIS cells with increasing doses of doxo and observed a dose-dependent increase in collagen degradation (Fig. S4A), suggesting that exposure to DNA damage induced a persistent increase in the collagen degradation capacity of DCIS cells.

This surprising result indicated that TREX1-dependent DNA damage in confined cells might favor cell invasion by increasing the collagen degradation capacity. We thus compared, in collagen assays, TREX1-proficient or -deficient (siTREX1 and TREX1 KO clones) DCIS cells, which had been confined for 2 h at either 2 μm or 10 μm height prior to embedding in the collagen gels. Once in the gel, the nuclear area of the cells, which had been confined at 2 μm or 10 μm, was comparable (Fig. S4B), showing no long-lasting nuclear deformation. TREX1-proficient cells that had been confined at 2 μm height showed a persistent, highly polarized morphology and increased speed in the collagen network (Fig. S4C, D and Movie S4). Consistent with our hypothesis, collagen degradation was augmented in TREX1-proficient cells that had been confined at 2 μm height, compared to 10 μm (Fig. 4D, E), but not in TREX1-depleted cells nor in TREX1 KO clones (Fig 4E-F). In agreement, DCIS cells overexpressing TREX1-WT showed augmented collagen degradation capacity following 2 μm confinement compared to non-overexpressing cells, whereas cells overexpressing TREX1-D18N displayed reduced collagen degradation compared to either non-overexpressing cells or cells overexpressing TREX1-WT (Fig. 4E). Importantly, doxo treatment induced collagen degradation in TREX1-depleted cells (Fig. S4E), confirming that TREX1 acts upstream of DNA damage to induce collagen degradation. Taken together, these experiments show that, upon strong confinement, TREX1-dependent DNA damage leads to a persistent increase in collagen degradation by breast tumor cells.

To further clarify how TREX1-mediated DNA damage induces collagen degradation upon confinement, cells confined at 2 μm height were treated with the MMP inhibitor, GM6001. This treatment inhibited collagen degradation, showing that confinement-induced collagen degradation depends on the activity of MMPs. Moreover, treatment of confined cells with the ATM inhibitor KU-55933 also reduced collagen degradation (Fig 4G), further confirming that confinement-induced collagen degradation requires the activation of the DNA damage response pathway. Finally, DNA damage in interphase is known to induce mitotic errors, chromosomal instability and the formation of micronuclei (^35^). These events lead to cGAS activation and an EMT-like phenotype (^35^). We thus wanted to test whether the cGAS signaling axis was underlying the increased collagen degradation following strong confinement of DCIS cells. Upon confinement at 2 μm height (Fig. S4F), or treatment with doxo (Fig. S4E), cGAS-depleted cells showed levels of collagen degradation comparable to control cells. This is consistent with the fact that induction of collagen degradation directly follows DNA damage induction by doxo or by confinement, while the cGAS-dependent mechanism takes place following at least one division event, and thus requires a long lag time following induction of DNA damage. Of note, we observed that cells do not divide while under confinement (Fig S3B) and we did not observe abnormal mitotic figures or micro-nuclei. Altogether, these results suggest that, in DCIS cells, strong confinement induces collagen degradation via a prolonged activation of the ATM-dependent DNA damage response pathway, independently of cGAS activation.

Our results so far showed that, in RPE1, MCF10A and DCIS cells, strong confinement leads to NE ruptures and induces chronic TREX1-dependent DNA damage. Contrary to RPE1 and MCF10A cells, confinement of DCIS cells does not induce senescence but leads to an ATM-dependent persistent increase in collagen degradation. Putting these results in the context of our initial observations of nuclear deformation and DNA damage in both human breast ductal carcinoma and in intraductal DCIS cell tumor xenografts (Fig 1A-F), we hypothesized that TREX1 might be responsible for DNA damage in these deformed nuclei, favoring the collagen degradation program, and thus promoting the transition from *in situ*-to-invasive carcinoma. To test this hypothesis, we injected either parental or TREX1 KO DCIS cells in the mammary ducts of mice and allowed tumors to develop for 6 to 8 weeks. Both parental- and TREX1 KO-injected mice formed intraductal *in situ* tumors at a similar rate (Fig. 5A). However, while in as early as 6 weeks, all the mice injected with parental DCIS cells showed invasive tumors based on whole-mount carmine and histological stainings, the formation of invasive tumors in TREX1 KO-injected mice remained greatly reduced for up to 8 weeks (Fig. 5B, C). Likewise, we observed a sharp decrease in the tumor area of TREX1 KO-injected mice compared to parental-injected mice (Fig. 5D). This phenotype was comparable to reduced invasive potential of Membrane Type I (MT1)-MMP (MMP14)-deficient DCIS cells (^17^). Importantly, the proliferation rates of parental DCIS and TREX1 KO clones were similar *in vitro* (Fig S5A). Next, 6-8 weeks tumors were labelled by immunohistochemistry with Ki67 to check whether cell proliferation was affected *in situ*, and with γH2AX to estimate the level of DNA damage *in situ*. Quantification of the fraction of Ki67 cells showed no significant difference in parental and TREX1 KO tumors (Fig 5E). Moreover, quantification of γH2AX-positive cells showed that in parental DCIS tumors, a fraction of the tumor slices analyzed showed a large number of γH2AX-positive cells, which were frequently found at the periphery of the tumors. In contrast, in TREX1 KO tumors, the number of γH2AX-positive cells remained low in all analyzed sections (Fig 5E, F, S5B and Movies S5 and S6). Overall, these results support our hypothesis and suggest that the transition from *in situ* to the invasive stage is favored in WT cells by TREX1-dependent DNA damage occurring in deformed nuclei as the tumor grows in the confined environment of the mammary ducts.

**Figure 5.**
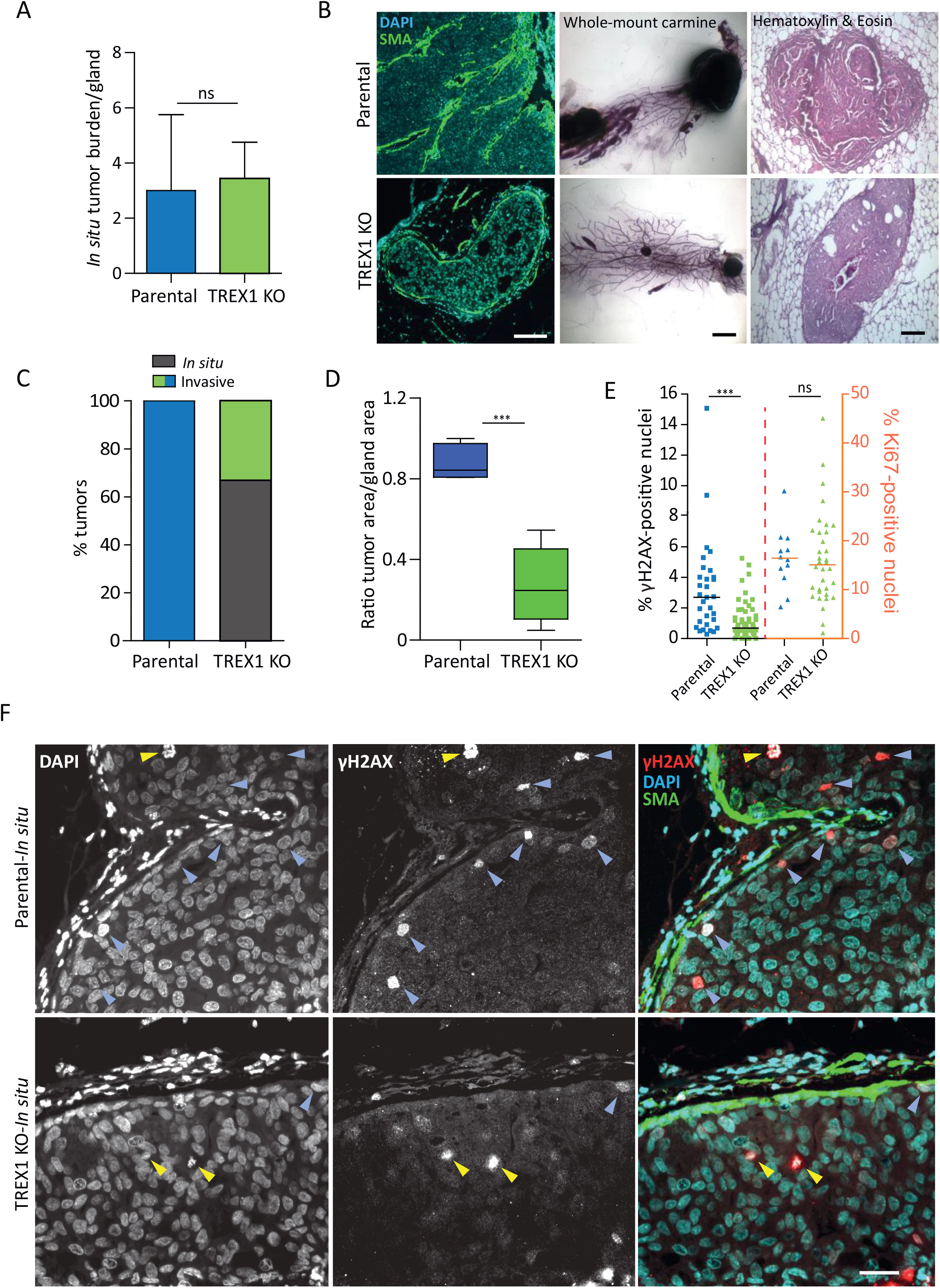
TREX1-dependent DNA damage promotes tumor invasion in mouse xenograft model of breast cancer progression. (A) Graph showing the rate of *in situ* tumor burden (period of 6-8 weeks post-intraductal injection). Data represents the mean ± SEM of 3 independent experiments where 20 cells per experiment per condition were analyzed. (B) Left panels: Immunofluorescence analysis of alpha-smooth muscle actin, SMA (green) and DAPI (cyan) in 6-8 weeks old mouse xenografts generated by intraductal injection of either parental DCIS cells or TREX1 KO clones. Bar, 100 μm. Middle panels: whole-mount carmine-stained 7 weeks old glands. Bar, 1 mm. Right panels: Hematoxylin & Eosin-stained 7 weeks old glands. Bar, 60 μm. (C) Graph showing the classification of the tumor xenograft stage (*in situ* or invasive) in the period of 6-8 weeks post-intraductal injection of either parental DCIS cells or TREX1 KO clones. Bars represent the mean ± SEM. (D) Graph: box and whisker plot showing the median value and 10-90 percentiles of the tumor area of *in situ* and invasive xenografts generated by DCIS parental or TREX1 KO clones (6-8 weeks post-intraductal injection). Bar, 100 μm. (A, C, D) Data represents a total of 7 glands scored for DCIS parental tumors and 14 glands for TREX1 KO tumors (TREX1 KO #1 +TREX1 KO #2 clones). (E) Scatter dot plots of γH2AX- and Ki67-positive nuclei (lines are median) 6-8 weeks post-intraductal injection of either parental DCIS cells or TREX1 KO clones. A total of 13 parental and 34 TREX1 KO (18 TREX1 KO #1 + 16 TREX1 KO #2) images were analyzed for Ki67 scoring (300 nuclei per image were scored); a total of 31 parental and 62 TREX1 KO (35 TREX1 KO #1 + 27 TREX1 KO #2) images were analyzed for γH2AX scoring (200 nuclei per image were scored). (F) Immunofluorescence analysis of γH2AX (red), apha-smooth muscle actin, SMA (green) and DAPI (cyan) in 6-8 weeks old mouse xenografts generated by intraductal injection of either parental DCIS cells or TREX1 KO clones. Blue arrowheads point to DNA damage-γH2AX-positive cells while yellow arrowheads point to mitotic cells. Bar, 40 μm. A total of 3 independent injection rounds to generate xenografts were performed. *P* values were calculated by unpaired Student’s *t*-test, ***P < 0.0001; ns = not significant

## Discussion

In this work, we have shown in different model systems (in several cell lines *in vitro*, in mouse tumor xenografts and in human samples of primary breast tumors) that a high level of DNA damage accompanies strong cellular and nuclear deformations. In our experimental models, ranging from confining micro-devices to mice, DNA damage is mostly dependent on the exonuclease TREX1. We also showed, *in vitro*, that TREX1-dependent DNA damage occurs only in nuclei deformed to the point of rupturing their NE, which provides a mechanistic explanation for the systematic correlation between strong nuclear deformations and DNA damage. To our knowledge, it suggests, for the first time, an enzymatic (possibly druggable) origin for DNA damage occurring in a variety of cell types and contexts in response to mechanical constraints (Fig. S6).

In addition, our data combining *in vitro* assays and mice xenograft tumors, suggest that this enzymatically driven chronic DNA damage in cells with deformed nuclei could lead to a senescence phenotype in normal cells and to a persistent collagen degradation in transformed cells, promoting cell invasion in collagen matrices and through the duct wall. Thus, our study establishes a mechanistic link between chronic DNA damage via the enzymatic activity of TREX1 and the progression of DCIS xenograft tumors in mice from the *in situ* to invasive stage.

TREX1 is a cytoplasmic exonuclease, mostly found to be associated with ER membranes (^36–39^), and shown to protect cells from accumulation of cytoplasmic DNA, either from viral or endogenous sources (^40–44^). Although TREX1 has already been proposed to act on nuclear DNA as well (^28,29,45^), possibly in conjunction with an endonuclease (^45^), to generate double strand breaks, there is so far no clear mechanism explaining how it would translocate from its cytoplasmic ER localization to act on the nuclear chromatin. ROS and ionic radiation have been shown to cause TREX1 nuclear translocation (^45–47^). Additionally, in the case of occurrence of NE ruptures, TREX1 could gain access to the nucleoplasm simply due to nucleo-cytoplasmic exchange, although this was never directly demonstrated. Our studies of TREX1-GFP localization in cells with NE ruptures show an increase in the nuclear localization of TREX1 as punctate foci, but the precise mechanisms of entry and action of TREX1 in the nucleus remain as open questions for future studies.

Similarly, to the best of our knowledge, there is no proposed mechanism establishing a direct link between DNA damage and collagen degradation. Our data suggest that, in the model systems we have studied, the ATM DNA damage response pathway is involved, rather than the cGAS-dependent inflammatory pathway (^35,48^), both for the senescence phenotype in RPE1 cells, and for the induction of the collagenolytic response in the senescence-deficient DCIS cell line. Of note, two recent studies pointed to a potential activation of an EMT-like phenotype downstream of ATM activation (^49,50^). Several ATM substrates have also been associated with lysosomal activity and thus potentially to collagen degradation (^51^). Our results suggest that upon chronic but sub-lethal levels of DNA damage, cells that have lost the normal ATM-dependent senescence-associated proliferation arrest, could promote an invasive behavior and drive the transition from *in situ* to invasive carcinoma stages.

A causal link between mechanical constraints, and in particular internal tumor pressure due to cell growth, and the activation of an invasive program has been put forward by several reports (^52^), but without a specific underlying molecular mechanism. Here we propose that tumor growth associated with the formation of peripheral motile strands of cells leads to strong deformation of the nucleus at the tumor edges. Nuclear deformation has been associated with several pathways (^53^), either triggering immediate cellular response, for example leading to an increase in cell contractility and motility (^54,55^), or to longer term responses associated with changes in gene expression and overall cell state or fate (^56^). Nuclear deformation was also shown to trigger YAP entry into the nucleus (^57^), and to trigger G1/S transition (^58^), thus potentially favoring cell proliferation. In contrast, a YAP-independent mechanotransduction pathway was recently proposed to drive breast cancer progression (^59^). Moreover, recent studies implicate YAP as a tumor suppressor during *in vivo* breast cancer progression (^60–62^). In our study, we show that a subset of cells, at the periphery of the tumor, can deform to the point of generating NE ruptures and DNA damage and that this phenomenon, which in some of the systems we studied is dependent on the nuclease TREX1, is responsible for an increase in cell invasiveness. How this phenomenon of strong nuclear deformation, NE ruptures and DNA damage interfaces with the YAP mechanotransduction pathway is an important question for future studies.

Beyond the fundamental interest of our findings, the fact that DNA damage is enzymatically generated in this context, suggests that cytoplasmic nucleases could be valuable new drug targets for cancer therapy. In this study, we pointed to TREX1, because its depletion in the cell lines we assayed was enough to abolish most of the DNA damage induced by strong confinement and to prevent senescence and invasive phenotypes, but other mechanisms could generate DNA damage in confined cells (^63^), which could be dependent or not on NE ruptures.

Our observation of DNA damage and deformed nuclei in primary breast tumor samples from patients suggests that confinement-associated DNA damage is likely to occur in the human disease. However, our experiments do not prove that TREX1 is the main driver of DNA damage in this context nor that DNA damage associated with nuclear deformation plays a role in the human disease progression. TREX1 has been described to promote pro- or anti-tumoral effects depending on different cellular and tumoral contexts (^64–66^). To get further insight into this question, we analyzed the association of TREX1 expression level in breast cancer with overall survival using public datasets. This analysis revealed that high level of TREX1 expression is significantly associated with reduced probability of survival (Fig S7). Notably, TREX1 is also implicated in modulating cGAS-dependent pro-tumorigenic inflammatory response (^40–42,44^). For exemple, TREX1 expression is reduced in fibroblasts undergoing replicative or oncogene-induced senescence, resulting in accumulation of DNA in cytosol and cGAS activation (^67^). Thus, targeting this endonuclease might be an effective therapeutic strategy to curb tumor progression, while promoting anti-tumoral immune response.

We and others have shown that the NE is a fragile barrier constantly being challenged in physiology and disease, and it ruptures at high frequency in deformed nuclei (^2,3^).The occurrence of DNA damage associated with nuclear deformations, possibly due to the enzymatic activity of cytoplasmic nucleases (depending on the cell type), is likely to be a common phenomenon. In normal tissues, this would trigger cellular senescence, contributing to developmental programs, tissue homeostasis, maybe serving as a cell-crowding sensor to regulate cell density, or even participating in tissue aging. This phenomenon could be exacerbated by mutations causing fragile nuclei, leading to degenerative diseases (^9,11,14,68–71^). In tumor cells, in which the senescence checkpoint has been lost, it would lead to a combination of aberrant invasive and proliferative behaviors, promoting tumor growth and invasion.

## Acknowledgements

We thank Titia de Lange (Rockefeller University, New York, USA) and John Maciejowski (Memorial Sloan Kettering Cancer Center, New York, USA) for providing the guideRNA to generate the TREX1 KO CRISPR cell lines. We also thank John Maciejowski and Alexis Lomakin for suggestions during the writing of this manuscript. We thank the flow cytometry and cell sorting facility and the animal facility at the Institut Curie (Paris, France) for provision of equipment and technical assistance. We thank Nicolas Carpi (Institut Curie, Paris, France) for lab management, plasmid amplifications and computer technical support. We thank Juan-Manuel Garcia Arcos (Institut Curie, Paris, France) for kindly drawing the model of this study (Figure S6). This work has also received the support of Institut Pierre-Gilles de Gennes-IPGG (Equipement d’Excellence, “Investissements d’avenir”, program ANR-10-EQPX-34). In addition, we thank the microscopy facility of IPGG for provision of equipment and technical assistance. Finally, we also thank the patients and their families who contributed tissue samples to these studies. S.A-G. was supported by a grant from the Laboratory of Excellence (LabEx) CelTisPhyBio ANR 11-LABX-0038; F.R. by a grant from Plan Cancer 2018 ‘Single Cells’ (19CS007-00). C.C. acknowledges financial support from the Fondation pour la Recherche Médicale (FDT20160435078). These studies were supported by the following grants: Associazione Italiana per la Ricerca sul Cancro (AIRC-IG#18621 to GS, AIRC-0IG#22145 to CT, and 5XMille #22759 to GS and CT); the Italian Ministry of University and Scientific Research (MIUR) to GS (PRIN: Progetti di Ricerca di Rilevante Interese Nazionale – Bando 2017#2017HWTP2K); the grants from Plan Cancer 2018 ‘Single Cells’ (19CS007-00) and Fondation ARC pour la Recherche contre le Cancer (PGA1 RF20170205408) and institutional funding from Institut Curie and Centre National de la Recherche Scientifique to P.C.; ERCadg CellO (FP7-IDEAS-ERC-321107) to JLV; LABEX DCBIOL (ANR-10-IDEX-0001-02 PSL* and ANR-11-LABX-0043), ANR ANR-17-CE15-0025-01 and ANR-18-CE92-0022-01, INSERM 19CS007-00, INCA PLBIO to N.M; INCA PLBIO 2019-1-PL BIO-07-ICR-1, INSERM Plan Cancer Single Cell grant 19CS007-00 to M.P.

## Author Contributions

G.P.F.N. designed and performed most of the experiments, analyzed and interpreted the data and helped M.P. writing the manuscript. S.A-G. and G.P.F.N. performed the 3D collagen degradation assays. S.A-G, A.Y., C.V., P.C. and J-L.V.designed the duct on chip device. S.A-G., F.R. and E.L. performed the mouse xenograft assays and S.A-G., F.R. and G.P.F.N. analyzed the data. M.Gr. conducted the molecular biology experiments necessary for the sequencing analysis of the TREX1 CRISPR clones and generated some of the stable cell lines. M.M. wrote Image J macros for data analysis and helped on data interpretation. C.C. analyzed and interpreted the data relative to cell cycle measurements. M.G. generated some of the stable cell lines and performed molecular cloning. C.L. provided images of mouse xenografts for analysis. V.C. and C.T. performed and interpreted the assays involving human primary tumors. G.S. designed, oversaw and interpreted the assays involving human primary tumors and edited the manuscript. P.C. designed, oversaw and interpreted the mouse xenograft assays and edited the manuscript. N.M. conceptualized the role of TREX1, helped designing and interpreting some of the experiments, supervised M.G. and edited the manuscript. G.P.F.N., S.A-G., M.P., P.C., N.M. and G.S. actively engaged in discussions throughout the study. M.P. supervised the study, designed the experiments, interpreted the data and wrote the manuscript.

## Supplementary informations

### Materials and Methods

#### Cell culture

The MCF10DCIS.com cell line was purchased from Asterand and maintained in DMEM-F12 medium with 5% horse serum. The MDA-MB-231 cell line was purchased from ATCC (ATCC HTB-26) and maintained in L-15 culture medium (Sigma-Aldrich, St Louis, MO, USA) with 2 mM glutamine (GIBCO, Cergy Pontoise, France) and 15% fetal bovine serum (GIBCO). MCF10A cells were culture in DMEM-F12 (Gibco) supplemented with EGF (20 ng/ml) (Peprotech #AF-100-15), hydrocortisone (0.5 μg/ml) (SIGMA #H0888), cholera toxin (100ng/ml) (SIGMA #C8052), Insulin (10μg/ml) (SIGMA #I9278), 5% horse serum and 1% penicillin and streptomycin (Lonza). RPE1 cells were grown in DMEM-F12 Glutamax medium (Gibco), supplemented with 10% FBS and 1% penicillin and streptavidin (Lonza). All cells were maintained at 37 °C in 5% CO_2_, with the exception of MDA-MB-231 cells, which were maintained at 37 °C in 1% CO_2_.

#### Lentiviral particles production in 293FT cells and Lentivector transductions

Lentiviral particles were produced as previously described from 293FT cells (^72^). Lentiviral viral particles and viral-like particles were produced by transfecting 1 μg of psPAX2 and 0.4 μg of pCMV-VSV-G together with 1.6 μg of a lentiviral vector plasmid per well of a 6-well plate. For cell transduction, 0.5×10^6^ cells were plated in a 6 well plate in 1ml and infected with 2 ml of fresh lentivector in the presence of 8 μg/ml of Protamine. The cells were then FACS-sorted by gating on the brightest GFP/mCherry-positive cells.

#### Constructs

The following constructs were produced in the lab of Nicolas Manel (Institut Curie, Paris, France): pTRIP-CMV-mCherry-TREX1 WT and D18N were cloned from pEGFP-C1-TREX1 (AddGene #27219) and pEGFP-C1-TREX1(D18N) (AddGene #27220) into pTRIP-CMV-mCherry-FLAG-cGAS E225A/D227A (^8^). pTRIP-SFFV-EGFP-FLAG-cGAS E225A/D227A and pTRIP-CMV-mCherry-FLAG-cGAS E225A/D227A are described elsewhere (^3,8^). pTRIP-SFFV-EGFP-53BP1 (amino acids 1224-1716 for isoform 1) was obtained by cloning from pTRIP-CMV-mCherry-53BP1 (^8^-Addgene #127658). The pIRES puro3 mCherry-Lifeact construct has been previously described (^73^).

#### Generation of TREX1 CRISPR clones

The guideRNA (gRNA) targeting TREX1 was kindly provided by the lab of Titia De Lange (Rockefeller University, New York, USA): gTREX1, 5’-GAGCCCCCCCACCTCTC-(PAM)-3’. gRNA plasmid was cotransfected into target cells with an hCas9 expression plasmid (Addgene) by nucleofection (Lonza apparatus). 1×10^6^ cells were mixed with electroporation buffer (freshly mixed 125 mM Na2HPO4, 12.5 mM KCl, 55 mM MgCl2 pH 7.75), 2 μg Cas9 plasmid, and 2 μg gRNA plasmid, transferred to an electroporation cuvette (BTX), and electroporated with program U-017 for RPE-1 cells or program D-023 for MCF10DCIS.com cells. Cells were then allowed to recover for 48 h before single cell FACs sorting into single wells of a 96 wells plate for single clone amplification and selection. Successful CRISPR/Cas9 editing was confirmed by both Western blotting of the amplified cell colonies and analysis of indel size (using the ICE analysis toolbox (https://www.synthego.com/products/bioinformatics/crispr-analysis).

#### Duct-on-a-chip assays

A 50 μm-high mold for the duct-on-chip chambers was made by standard photolithography, using SU-8 2050 photoresist (Microchem) spin-coated on a silicon wafer (Neyco), which was exposed to UV light through a chromium mask (MB Whitaker & Associates) created with a micro pattern generator μPG 101 (Heidelberg Instruments). The chamber was made by molding polydimethylsiloxane (PDMS, RTV615, GE) and bonded on a FluoroDish. Unlabeled type I collagen at 2.4 mg/ml (BD Biosciences, Cat. # 354236) was injected in the “collagen chamber” of the microfabricated duct-on-a-chip and incubated for 1h at 37°C. In the meantime cells were ressuspended in matrigel (SIGMA #E1270) at a concentration of 0.7×10^6^ cells/μl and then 3 μl were injected in the “cells chamber”. Images were acquired at the desired time points after injection to assess cell invasion into the “collagen chamber”.

#### Human primary tumor samples

Human breast cancer tissue samples were collected according to the Helsinki Declaration and the study was approved by the University of Palermo Ethical Review Board (approval number 09/2018). The cases were classified according to the World Health Organization classification criteria of the tumors of the breast by trained pathologist, author of this article (Claudio Tripodo). Sections 2.5/3 micron-thick were cut from paraffin blocks, dried, de-waxed and rehydrated. The antigen unmasking technique was performed using Target Retrieval Solutions pH6 at 98°C for 30 min. After neutralization of the endogenous peroxidase with 3% H_2_O_2_ and Fc blocking by a specific protein block, double-marker immunohistochemistry was carried out by incubation for 90 min at RT with the primary antibodies RAB55a (Abcam, 1:100 pH6) and γH2AX (Abcam, 1:1000 pH6). Staining was revealed using Novolink Polymer Detection Systems (Novocastra) and SuperSensitive Link-Label IHC Detection System Alkaline Phosphatase (Biogenex). DAB (3,3’-diaminobenzidine) and Vulcan Fast Red were used as substrate chromogens. The slides were counterstained with Harris hematoxylin (Novocastra) to reveal nuclei. All the sections were analyzed under a Zeiss AXIO Scope.A1 microscope (Zeiss, Germany) and microphotographs were collected using a Zeiss Axiocam 503 Color digital camera using the Zen2 imaging software. Double immunohistochemistry was performed on 7 breast cancer tissues, but taking into account also the double IF RAB5A/gH2aX, the number of cases increases to 15.

#### Intraductal transplantation method

The intraductal xenograft model was carried out as previously described (^17,24^). Briefly, 5×10^4^ MCF10DCIS.com cells in 2 μl PBS were injected into the primary duct through the nipples of both mammary inguinal glands #4 of 8-10 weeks-old virgin female SCID mice. Mice were sacrificed from 5 to 8 weeks after injection by cervical dislocation. Immediately after being euthanized, mammary glands were excised and processed for further study (including wholemount and histological and IHC staining on sections). Animal care and use for this study were performed in accordance with the recommendations of the European Community (2010/63/UE) for the care and use of laboratory animals. All animal experimental procedures were approved by ethics committee of Institut Curie CEEA-IC #118 (Authorization APAFiS #24649-2020031220398710-v1 given by National Authority) in compliance with international guidelines.

#### Histological and immunofluorescence analysis of mouse tissue sections

Whole-mount carmine and hematoxylin and eosin staining were performed as described (^74^). To retrieve antigens on paraffin-embedded tissue samples, sections were incubated for 20 min in 10 mM sodium citrate buffer, pH 6.0 at 90°C. Then, after 1 h incubation in 5% fetal calf serum, sections were incubated overnight with diluted primary antibodies, washed and further incubated for 2h at room temperature with appropriate secondary antibodies.

#### Indirect immunofluorescence microscopy

Samples were fixed with 4% paraformaldehyde, permeabilized with 0.1% Triton X-100, and then incubated for 1 h at room temperature with indicated antibodies. Following three washes with PBS samples were incubated with the appropriate secondary antibodies for 1 h at room temperature. Samples were again washed three times with PBS and mounted with Fluormount G (Molecular probes).

#### 3D type I collagen degradation assay

For collagen degradation assessment followed by immunofluorescence analysis, glass bottom dishes (MatTek Corporation) were layered with 15 μl of a solution of 5 mg/ml unlabeled type I collagen (IBIDI Cat. # 50201) mixed with 1/20-40 volume of Alexa Fluor 647-labeled collagen. Polymerization was induced at 37°C for 3 min and complete medium was added. Cells were seeded onto the polymerized collagen layer and incubated for 1h at 37 °C. The medium was then gently removed and two drops of a mix of Alexa Fluor 647-labeled type I collagen (10% final) and unlabeled type I collagen at 2.4 mg/ml (BD Biosciences, Cat. # 354236) were added on top of the cells (top layer). After polymerization at 37°C for 1.5 hrs, 1 ml of medium was added to the MatTek dishes and cells were incubated for XXX hrs at 37°C. After fixation for 30 min at 37 °C in 4% paraformaldehyde in PBS, samples were incubated with anti-Col1-3/4C antibodies for 2 h at 4 °C. After extensive washes, samples were counterstained with Cy3-conjugated anti-rabbit IgG antibodies, Phalloidin-Alexa488 to visualize cell shape and mounted in DAPI. Image acqui-sition was performed with an A1R Nikon confocal microscope with a 40X NA 1.3 oil objective using high 455 sensitivity GaASP PMT detector and a 595 ± 50 nm band-pass filter. Quantification of degradation spots was performed as described 15. Briefly, maximal projection of 10 optical sections with 2 μm interval from confocal microscope z-stacks (20 μm depth) were preprocessed by a Laplacian-of-Gaussian filter using a homemade ImageJ macro 15. Detected spots were then counted and saved for visual verification. No manual correction was done. Degradation index was the number of degradation spots divided by the number of cells present in the field, normalized to the degradation index of control cells set to 100 (^75^).

#### Transfection procedure and siRNA oligonucleotides

For RNA interference experiments, cells were transfected with siRNA oligonucleotides (Dharmacon) using Lipofectamine^®^ RNAiMAX reagent (Invitrogen) (RPE1 cells), or with Lullaby (OZ Biosciences, France) (MCF10A cells, MCF10DCIS.com cells and MDA-MB-231 cells) according to manufacturer’s protocol. The following SMARTpool siRNAs were used: human TREX1 (Dharmacon, cat. # L-013239-02-0005), human MB21D1 (cGAS) (Dharmacon, cat. # L-015607-02-0005) and validated non-targeting siRNAs (Dharmacon, cat. # D-001810-10-20). Cells were analyzed 72 h post-transfection using standard Western blot or immunofluorescent analysis protocols. SMARTpool siRNAs sequences:

ON-TARGETplus SMARTpool TREX1 siRNAs (siTREX1): 5′-GCCACAACCAGGAACACUA-3′, 5′-GCGCAUGGGCGUCAAUGUU-3′, 5′-CAGAACACGGCCCAAGGAA-3′, 5′-UGUCACAACCACUUGCACAC-3′; ON-TARGETplus SMARTpool MB21D1 siRNAs (sicGAS) 5′-GAAGAAACAUGGCGGCUAU-3′, 5′-AGGAAGCAACUACGACUAA-3′, 5′-AGAACUAGAGUCACCCUAA-3′, 5′-CCAAGAAGGCCUGCGCAUU-3′; ON-TARGETplus non-targeting SMARTpool siRNAs (siControl).

#### Drug treatments

The following pharmacological inhibitors and chemical compounds were used: Etoposide (DNA damaging agent, topoisomerase-II inhibitor; Cell Signaling, #2200), Doxorubicin (DNA damaging agent, topoisomerase-II inhibitor; Cell Signaling, #5927), GM6001 (pan-MMP inhibitor); Millipore, # CC1010), KU-55933 (ATM kinase inhibitor; Abcam, #120637).

#### Western blotting

Cells were collected and resuspended in Laemmeli buffer. Proteins were separated using sodium dodecyl sulfate polyacrylamide gel electrophoresis (SDS–PAGE) and transferred onto nitrocellulose membranes. After incubation with primary antibody the membrane was incubated with and goat horseradish peroxidase-conjugated anti-rabbit IgG (Sigma A0545, 1/10 000) or anti-mouse IgG antibodies (Jackson ImmunoResearch 61871, 1/10 000).

#### Microfabrication-based confinement

##### Confinement using the “6 well confiner”

To obtain large quantities of confined cells for cell population or biochemical studies, cell confinement was performed using a version of the cell confiner adapted to multi-well plates (^76^). To make the PDMS microspacers (micropillars) at the desired height, 12 mm glass coverslips were plasma treated and then placed on top of a PDMS mixture (10/1 w/w PDMS A / crosslinker B) on the wafer molds (containing holes/micropillars, fabricated following standard photolithography procedures). The height of the micropillars determines the height for spatial confinement of cells between the coverslip and the substrate. The surface of the confining side was always treated with non-adhesive pLL-PEG (SuSoS, PLL(20)-g[3.5]-PEG(2)). After baking at 95°C for 15 min, coverslips with PDMS pillars were carefully removed from the wafers under isopropanol. They were then cleaned with isopropanol, well-dried, treated with plasma for 1 min, and treated with 0.5 mg/mL pLL-PEG in 10 mM pH 7.4 HEPES buffer for 1h at room temperature. Coverslips with PDMS pillars were rinsed and incubated in medium for at least 2 h before confining the cells. The modified cover-lid of a multi-well plate was used to apply confining slides to cells. In this case, large PDMS pillars were stuck on the cover-lid of the multi-well plate to hold the confining slides containing a layer of microfabricated micropillars. These large PDMS pillars push the confining slides from the top of the plastic 6 well cover-lid to confine the cells in 6 well glass/plastic bottom plates. The process of fabrication for these large pillars attached to the 6 well plate lid is as follows: the large PDMS pillars were fabricated by pouring a PDMS mixture (A:B = 35:1) into a custom-made metallic mold, removing bubbles under vacuum, then baking overnight at 80°C, and getting the pillars out of the mold with the help of small amount of isopropanol.

#### Antibodies

For immunofluorescence and IHC analysis on tissue sections, the following primary antibodies were used: Ku70 (clone EPR4026, Abcam, #ab108604, 1/100), Ki-67 (clone MIB-1, Dako, 1/100), αSMA (clone 1A4, Dako, 1/100), Col1-3/4C (collagenase-cleaved ¾ fragment of collagen I, ImmunoGlobe GmbH, #0217-050). Nuclei with 4’,6-diamidino-2-phenylindole (1 μg/ml, Sigma, Saint-Quentin Fallavier, France). TREX1 (Cell Signaling, #12215), TREX1 (Clone E-6, Santa Cruz Biotech, #sc-271870), phospho-Histone H2A.X (Ser139) (Clone JBW301, Millipore #05-636), RIF1 (Bethyl laboratories, #A300-569A), cGAS (Clone D1D3G, Cell Signaling, #15102), p21 (Clone EPR3993, Abcam, #ab109199), HP1γ (Clone 14D3.1, Millipore, #MABE656), Tubulin (Clone DM1A, SIGMA, #T9026), GAPDH (Clone 14C10, Cell Signaling, #2118), laminB1 (Clone A-11, Santa Cruz Biotech, #sc-377000).

Secondary antibodies anti-mouse IgG-Cy3 or -Cy2 (Jackson ImmunoResearch, Montluçon, France, 63578, 1/500) and anti-rabbit-IgG-Cy3 or -Cy2 (Life Technologies, Cergy Pontoise, France, A-11034, 1/500) were used.

#### Live-cell imaging

Time-lapse recordings were acquired with 20x (NA 0.75) or 40x (NA 0.85) dry objectives using either an Eclipse Ti inverted microscope (Nikon) equipped with a Coolsnap HQ2 camera (Photometrics) or with a spinning-disc confocal microscope with a Yokogawa CSU-X1 spinning-disc head on a DMI-8 Leica inverted microscope equipped with a Hamamatsu OrcaFlash 4.0 Camera, a NanoScanZ piezo focusing stage (Prior Scientific) and a motorized scanning stage (Marzhauser). Both microscopes were controlled by MetaMorph software (Molecular Devices). All microscopes were equipped with an on-stage incubation chamber which maintained the temperature at 37°C and CO_2_ concentration at 5% at all times. Image analysis was performed using ImageJ/Fiji software (NIH, http://rsb.info.nih.gov/ij/index.html) or MetaMorph software (Universal Imaging).

#### Statistics and reproducibility of experiments

Unless stated otherwise, statistical significance was determined by one-tailed unpaired Student’s *t*-test after confirming that the data met appropriate assumptions (normality, homogenous variance and independent sampling). Statistical data are presented as median or mean ± either SEM or SD. Sample size (n) and p-value are specified in the figure legends. Samples in most cases were defined as the number of cells counted/examined within multiple different fields of view on the same dish/slide, and thus represent data from a single sample within a single experiment, that are representative of at least three additional independently conducted experiments.

## Supplementary Figures Legends

**Figure S1.**
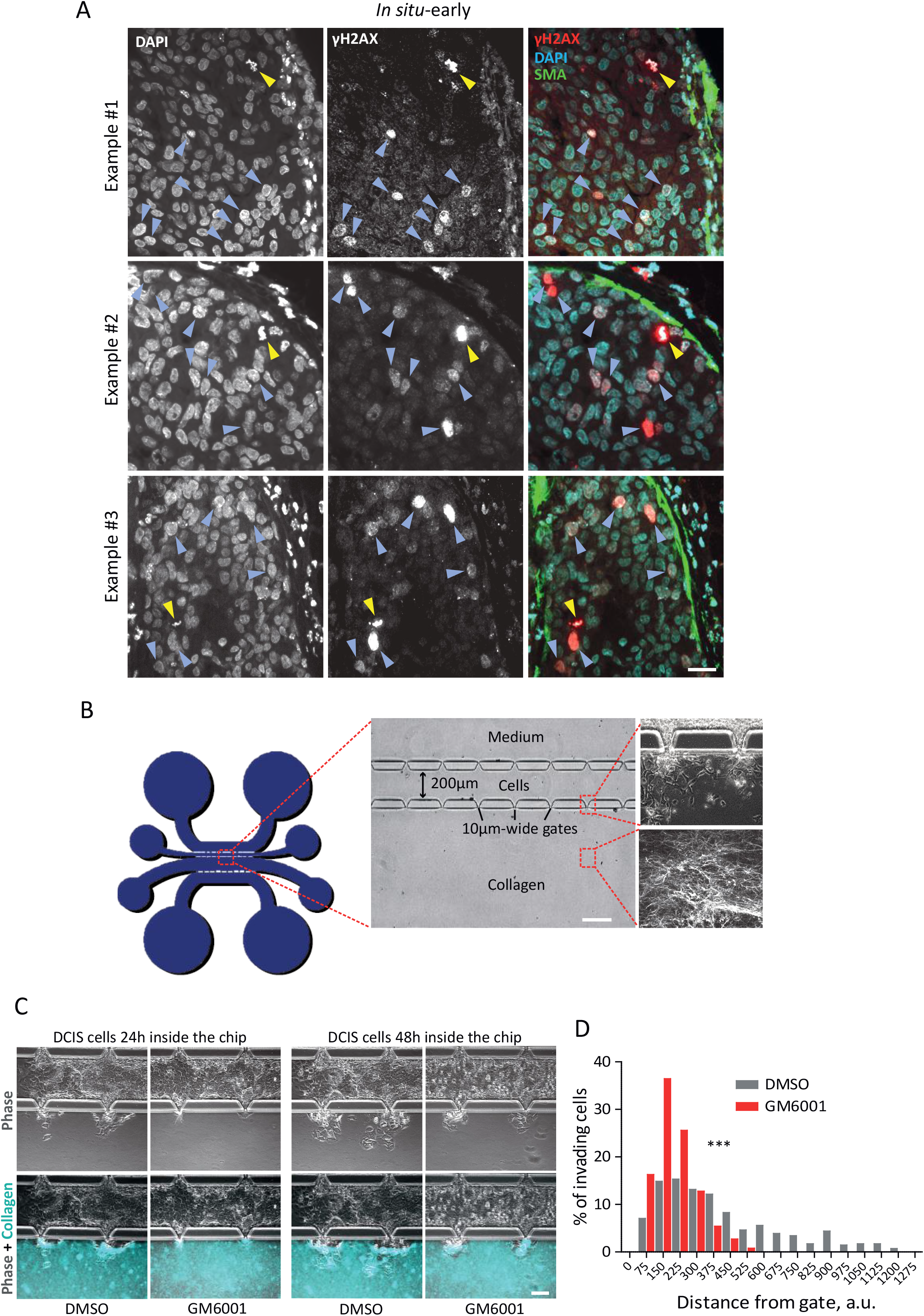
The “duct-on-chip” device: reconstitution of the mammary duct using microfabrication. (A) Immunofluorescence analysis of mouse xenografts at i*n situ*-early stage generated by intraductal injection of DCIS cells (6-8 weeks post-intraductal injection). Alpha-smooth muscle actin, SMA (green), human-specific KU70 (cyan) and γH2AX (red). Blue arrowheads point to DNA damage-γH2AX-positive cells while yellow arrowheads point to mitotic cells. Bar, 40 μm. (B) Diagram illustrating the duct-on-chip device. A mixture of cells and matrigel is injected in the “cell chamber” and type I collagen is injected in the collagen chamber. Subsequently, culture medium is injected in the “medium chamber”. Note that cells invade towards the collagen chamber through 10 μm wide gates that recapitulate the breaches in the myoepithelial layer during *in vivo* of cell invasion. Bar, 80 μm. (C) DCIS cells were injected in the duct-on-chip in the presence of DMSO (vehicle) or MMP pan inhibitor (GM6001, 40 μM). Cell invasion was measured as distance from the gates. Graph: frequency distribution of invading cells as a function of the distance invaded from the gate. Data represents 3 independent experiments where cells invading from 10 random gates (at 20x magnification) were analyzed per condition per experiment. Bar, 60 μm. *P* values were calculated by unpaired Student’s *t*-test, ***P < 0.0001.

**Figure S2.**
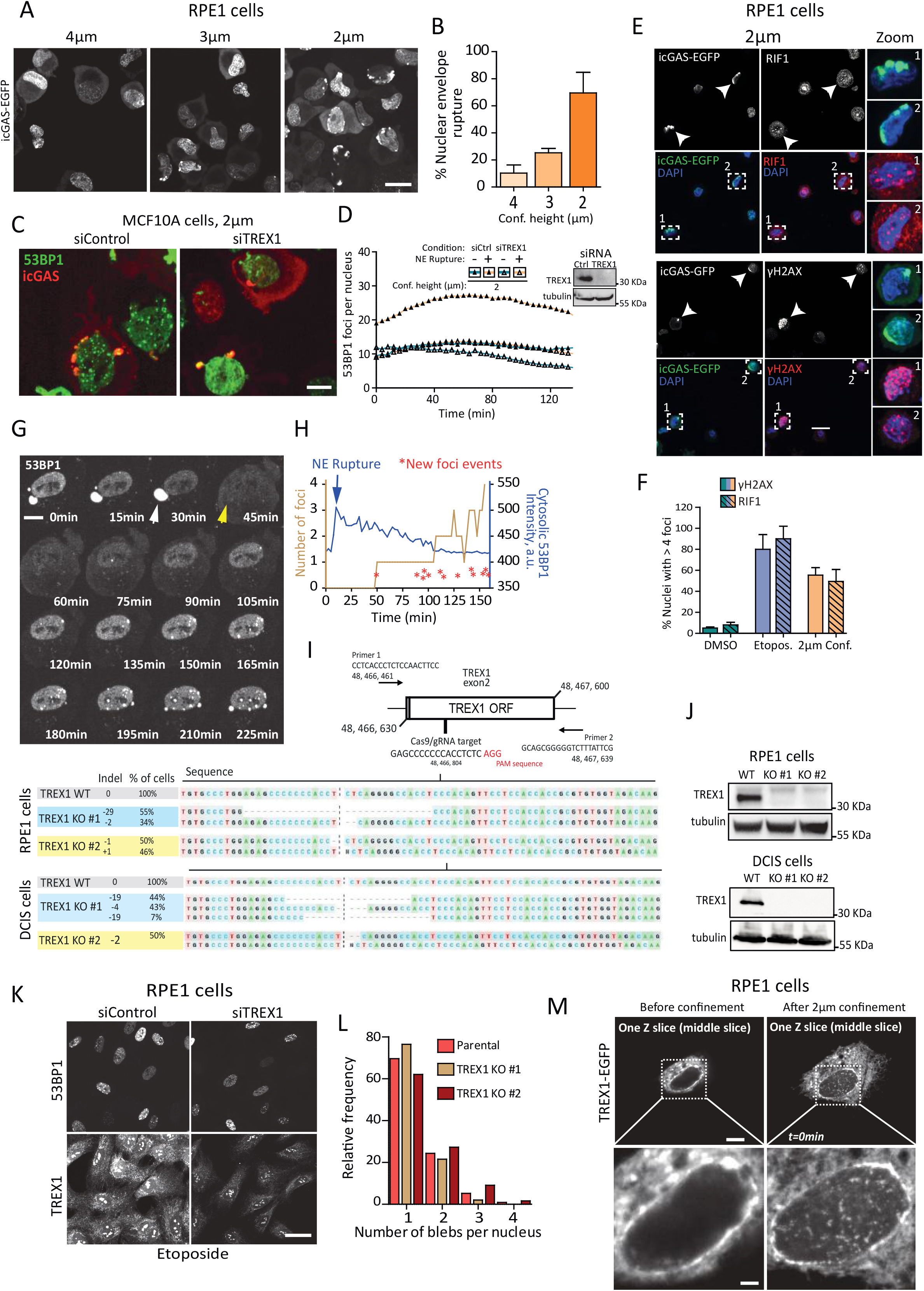
Characterization of nuclear envelope rupture under different confinement heights and of DNA damage following nuclear envelope rupture. (A) RPE1 cells stably expressing catalytically inactive cGAS-EGFP were confined at the indicated heights and images were acquired immediately after (and while cells were under confinement). Bar, 15 μm. (B) Quantification of NE rupture events as assessed by cGAS perinuclear localization. Data represents the mean ± SD of 3 independent experiments where 50 cells per experiment per height were analyzed. (C) MCF10A cells stably expressing 53BP1-EGFP and catalytically inactive cGAS-mCherry were transiently depleted for TREX1 with siRNA and 48 h later cells were confined at 2 μm. (D) Quantification of DNA damage levels during 2 μm confinement as assessed by the number of 53BP1 foci in siControl and siTREX1 cells displaying or not NE rupture. Western blot shows TREX1 depletion 48 h post-knockdown; tubulin is the loading control. Bar, 10 μm. Data represents the mean of 3 independent experiments where 20 cells per experiment per condition were analyzed. (E) RPE1 cells stably expressing catalytically inactive cGAS-EGFP were confined at 2 μm for 2 h; subsequently, the confinement lid was removed, cells were harvested, replated for 30 minutes and fixed for immunostaining with the DNA damage markers RIF1 and γH2AX (red) and DAPI (blue). Arrowheads point to cells with NE rupture, which are also positive for the DNA damage markers. Bar, 20 μm. (F) Quantification of DNA damage foci at the indicated conditions (DMSO, etoposide 25 μM and 2 μm confinement). Images are representative of 2 independent experiments where 30 cells per experiment per condition were analyzed. Bars represent the mean ± SD. (G) RPE1 cells stably expressing 53BP1-EGFP were confined at 2 μm and imaged under spinning disc microscopy. White arrowhead points to a NE bleb while the yellow arrowhead points to a bleb bursting event that is followed by the appearance of DNA damage foci. (H) Graphs showing the absolute number of 53BP1 foci and the events of appearance of new 53BP1 foci (red asterisks) following a bleb bursting event (assessed by an increase in the cytosolic intensity of the probe 53BP1-EGFP, which leaks out of the nucleus upon NE rupture). Blue arrow indicates the instant of NE rupture/bleb bursting. (I) Diagrams illustrating the Cas9/gRNA targeting sequence in the *TREX1* gene and the primers used for the sequencing of the TREX1 KO clones generated by CRISPR technology for RPE1 and DCIS cells. (J) Western blots of WT and TREX1 KO clones; tubulin is a loading control. (K) RPE1 cells stably expressing 53BP1-EGFP were transiently depleted for TREX1 using siRNA and treated with etoposide (25 μM) for 2 h. Subsequently cells fixed and stained for endogenous TREX1. Bar, 25 μm. (L) Frequency distribution of the number of nuclear blebs in RPE1 TREX1 KO clones and in parental RPE1 cells. Data represents 3 independent experiments where 60 cells per experiment were analyzed. (M) RPE1 cells transiently transfected with TREX1-EGFP WT were confined at 2 μm using a pressure-controlled dynamic confiner. Images represent a single plane Z slice through the middle section of the nucleus. Bars, 5 μm and 1 μm (magnification). Images are representative of 2 independent experiments.

**Figure S3.**
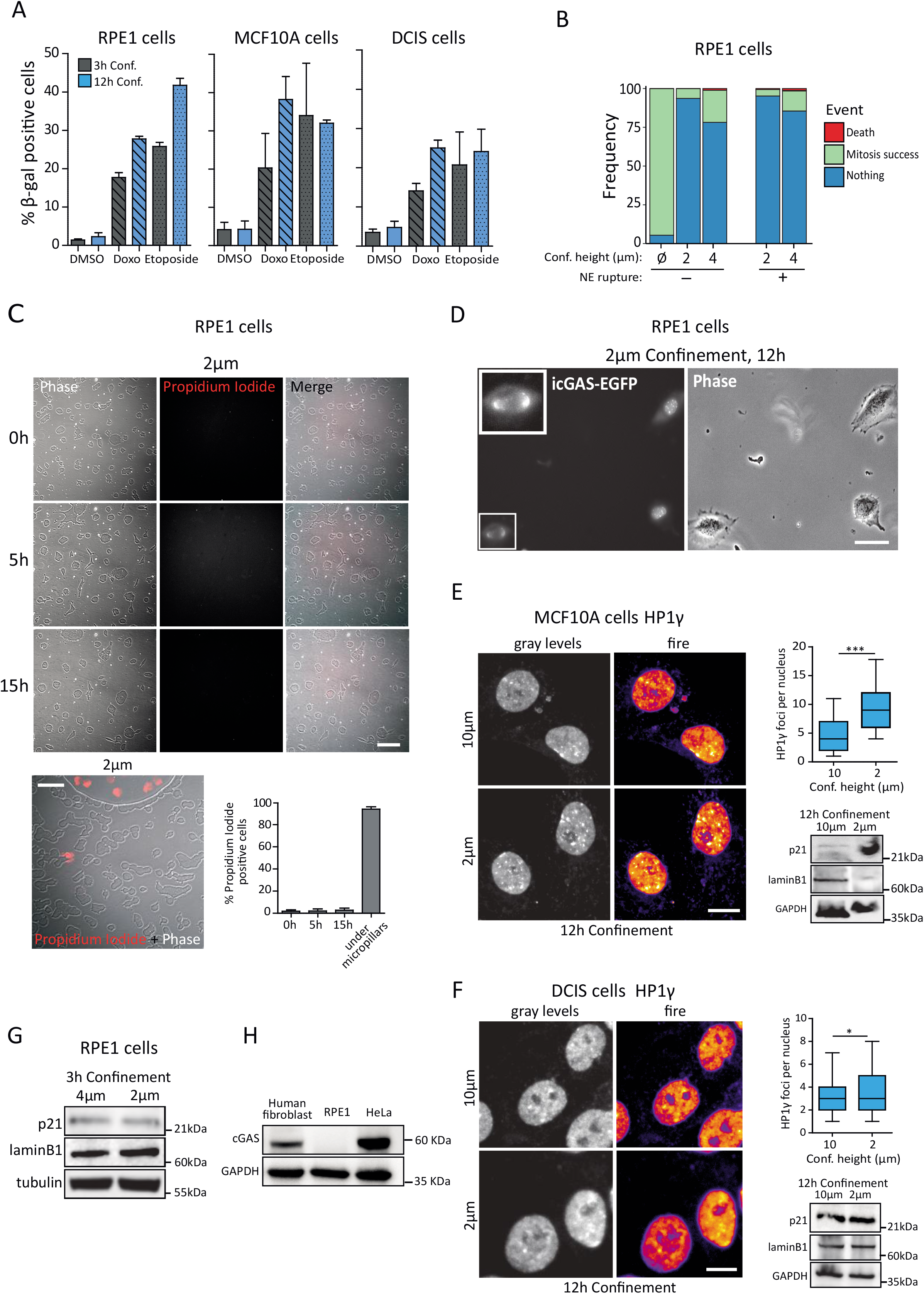
Long-term, strong confinement causes cell senescence in non-transformed cells but not in cancer cells. (A) RPE1, MCF10A and DCIS cells were treated with doxorubicin (40 nM), etoposide (25 μM) or DMSO (vehicle) for the indicated lengths of time. Cells were then fixed and processed for β-gal staining. The percentage of β-gal-positive cells was plotted. Graph: data represents mean ± SD of 3 independent experiments where 200 cells were scored per condition per experiment. (B) Frequency distribution (percentage) of the designated categories for cells under the indicated confinement heights. Data represents 3 independent experiments where 37 non-confined cells, 40 confined cells with non-ruptured nuclei and 60 confined cells with ruptured nuclei were scored per condition per experiment. (C) RPE1 cells were confined at 2 μm in the presence of propidium iodide and imaged overnight. Graph: data represents the mean ± SD of 2 independent experiments where 5 random fields (100 cells scored per field) were analyzed per time-point per experiment. Bar, 50μm. (D) Epifluorescence and phase images of RPE1 cells stably expressing catalytically inactive cGAS-EGFP harvested from 2 μm confinement and replated for cell cycle duration measurement over a 72 h period. Bar, 25 μm. Inset shows a cell with ruptured NE, as evidenced by cGAS perinuclear accumulation. (E) MCF10A cells or (F) DCIS cells were confined for 12 h at 10 or 2 μm, harvested from confinement and replated for 72 h before fixation for immunostaining with heterochromatin foci (HP1γ) or lysis for western blot analysis of lamin B1 and p21. GAPDH is the loading control; western blot images are representative of 2 independent experiments. Graph: box and whisker plot showing the median value and 10-90 percentiles of the number of HP1γ foci per nucleus. Data represents 3 independent experiments where 100 cells per condition per experiment were analyzed. Bars, 10 μm. (G) RPE1 cells were confined for 12 h at 4 or 2 μm; cells were then harvested from confinement and replated for 72 h before lysis for western blot analysis of lamin B1 and p21. Tubulin is the loading control; western blot images are representative of 2 independent experiments. (H) Western blot analysis of whole cell extracts of human fibroblasts, RPE1 cells and HeLa cells. GAPDH is the loading control. *P* values were calculated by unpaired Student’s *t*-test, ***P < 0.0001; *P < 0.05; ns = not significant.

**Figure S4.**
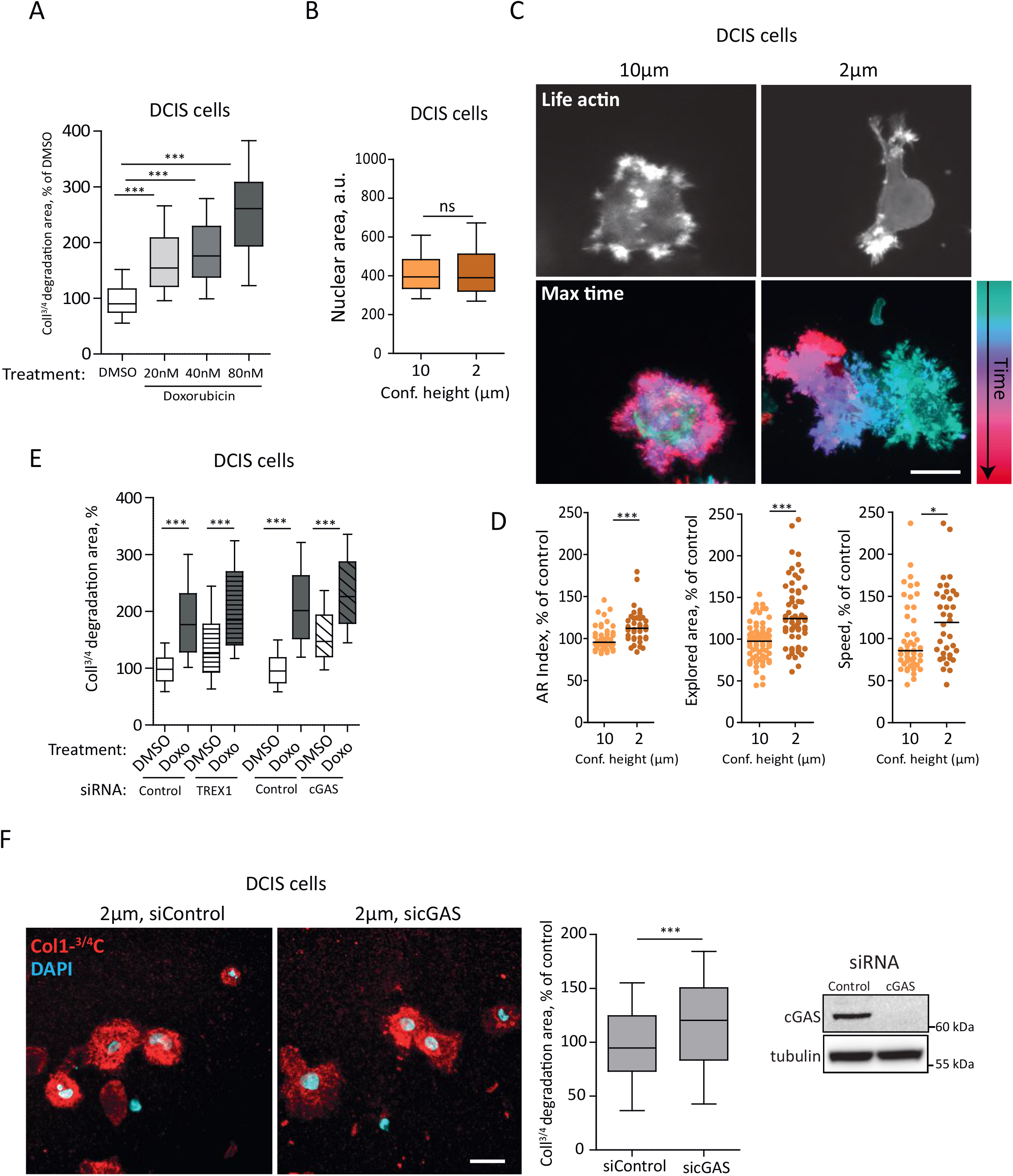
Pharmacologically induced DNA damage promotes collagen degradation independently of TREX1 and cGAS and strong confinement induces cell polarization and collagen degradation in a cGAS-independent fashion. (A) DCIS cells were treated for 12 h with the indicated concentrations of doxorubicin or DMSO (vehicle). Following the incubation period, cells were harvested and embedded in 3D type I collagen for 12 h to assess their collagen degradation activity by immunostaining with an antibody that specifically recognizes the collagenase-cleaved ¾ fragment of collagen I. Graph: box and whisker plot showing the median value and 10-90 percentiles of the collagen degradation area. Data represents 3 independent experiments where 50 cells per condition per experiment were analyzed. (B) Quantification of nuclear area of DCIS cells embedded into 3D type I collagen after being confinedfor 2h at the indicated heights. Graph: box and whisker plot showing the median value and 10-90 percentiles of nuclear projected area. Data represents 3 independent experiments where 70 cells per condition per experiment were analyzed. (C) DCIS cells stably expressing life actin-EGFP were confined for 2 h at the indicated heights. Following the confinement period, cells were harvested and embedded in 3D type I collagen and imaged for 15 h. Bar, 20 μm. (D) Scatter dot plots of different measured parameters while cells where embedded into 3D type I collagen (lines are median). Data represents 2 independent experiments where 25 cells per condition per experiment were analyzed. (E) DCIS cells were transiently depleted for TREX1 or cGAS using siRNA and 48 h later cells were treated for 12 h with doxorubicin (Doxo, 80 nM) or DMSO (vehicle). Following the treatment period, cells were harvested and embedded in 3D type I collagen for 12 h to assess their collagen degradation activity by immunostaining with an antibody that specifically recognizes the collagenase-cleaved ¾ fragment of collagen I. Graph: box and whisker plot showing the median value and 10-90 percentiles of the collagen degradation area. Data represents 3 independent experiments where 40 cells per condition per experiment were analyzed. (F) DCIS cells were transiently depleted for cGAS using siRNA and 48 h later cells were confined at 2 μm for 2 h. Following the confinement period, cells were harvested and embedded in 3D type I collagen for 12 h to assess their collagen degradation activity by immunostaining with an antibody that specifically recognizes the collagenase-cleaved ¾ fragment of collagen I (red); DAPI (cyan). Bar, 20 μm. Graph: box and whisker plot showing the median value and 10-90 percentiles of the collagen degradation area. Data represents 3 independent experiments where 60 cells per condition per experiment were analyzed. Far right: western blot analysis of cGAS depletion (48 h post-knockdown); tubulin is a loading control. *P* values were calculated by unpaired Student’s *t*-test, ***P < 0.0001; *P < 0.05; ns = not significant.

**Figure S5.**
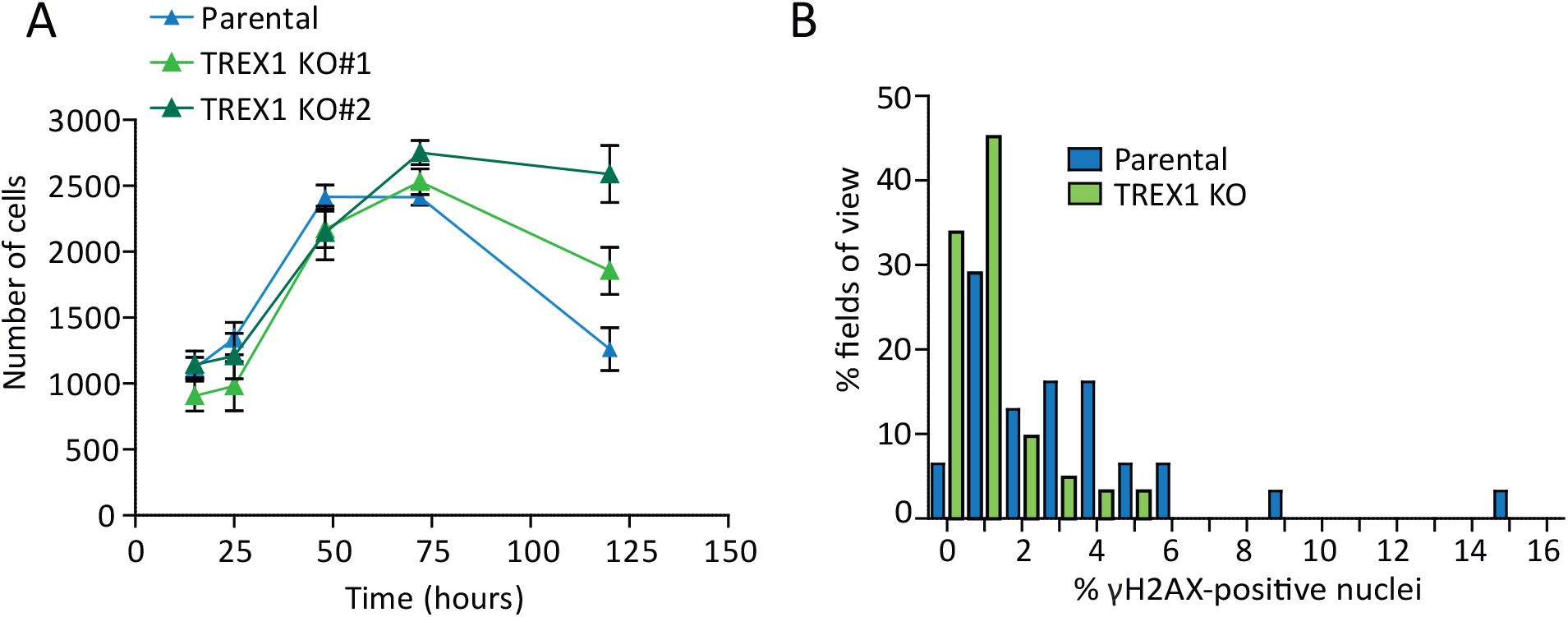
TREX1 KO clones display normal proliferation *in vitro*. (A) Proliferation curve of DCIS parental, TREX1 KO#1 and TREX1 KO#2 clones. Graph: data represents the mean ± SEM of 25 random fields scored at 20x magnification. (B) Frequency distribution of the percentage of γH2AX-positive nuclei in DCIS parental *in situ* xenografts and TREX1 KO *in situ* xenografts 6-8 weeks post-intraductal injection. A total of 31 parental and 62 TREX1 KO (35 TREX1 KO #1 + 27 TREX1 KO #2) images were analyzed for γH2AX scoring (200 nuclei per image were scored).

**Figure S6.**
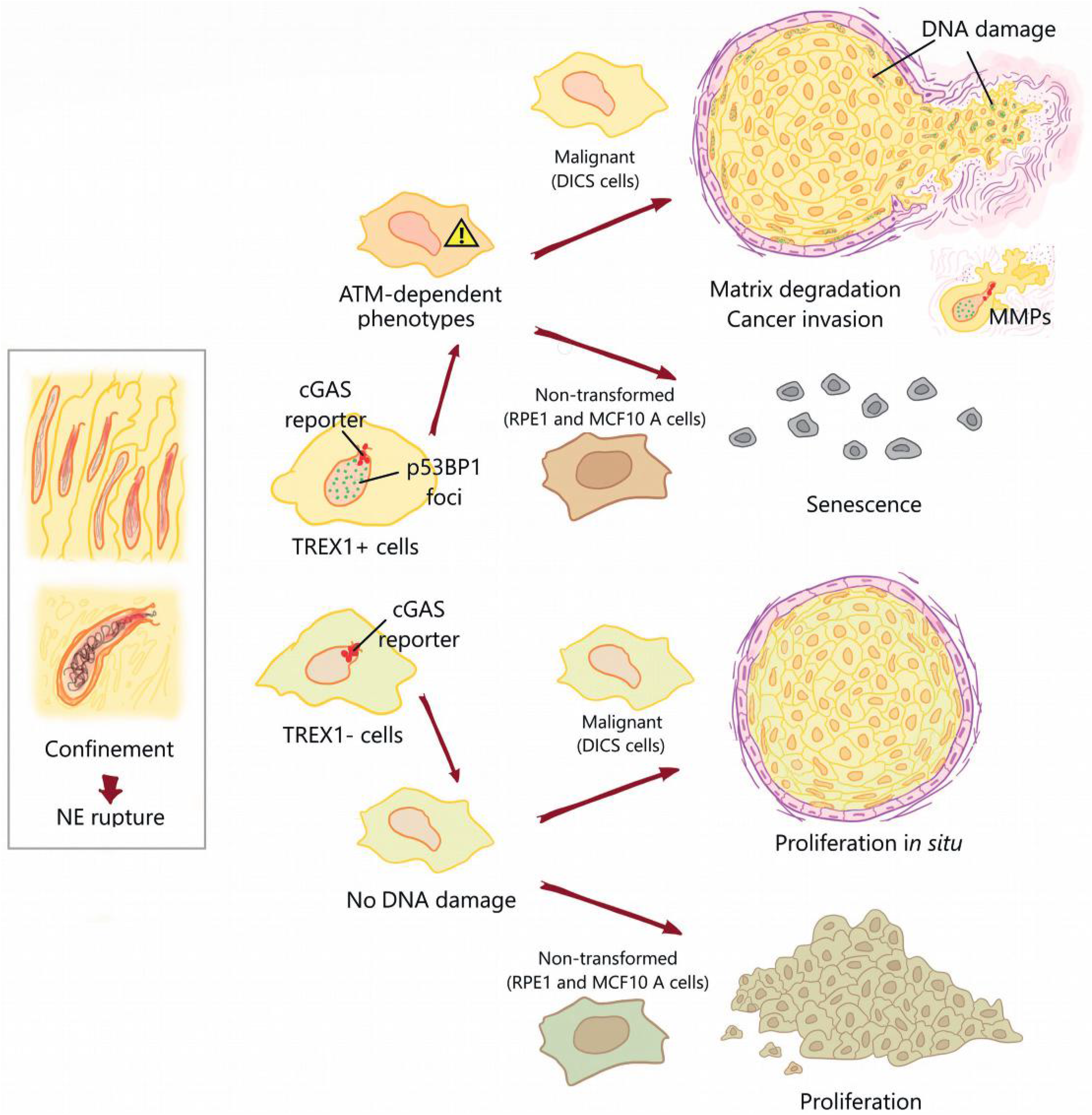
Schematic model of nuclear deformation, rupture and TREX1-dependent DNA damage in confining 3D microenvironments. The model depicts the consequences of strong nuclear envelope deformations that cells experience when exploring and growing in dense *in vivo* environments. Upon nuclear envelope rupture (cGAS perinuclear accumulation, red), the exposed nuclear DNA becomes a substrate for the nuclease TREX1, which generates DNA damage (53BP1 foci, green). Repeated events of nuclear envelope rupture will cause a chronic DNA damage response (ATM activation) and will have two outcomes: in non-transformed/healthy cells it triggers cell senescence while in transformed cells it promotes matrix degradation and cancer invasion. In contrast, in TREX1 KO cells the absence of DNA damage upon nuclear envelope rupture prevents the activation of the DNA damage response in transformed cells, thus preventing cancer invasion.

**Figure S7.**
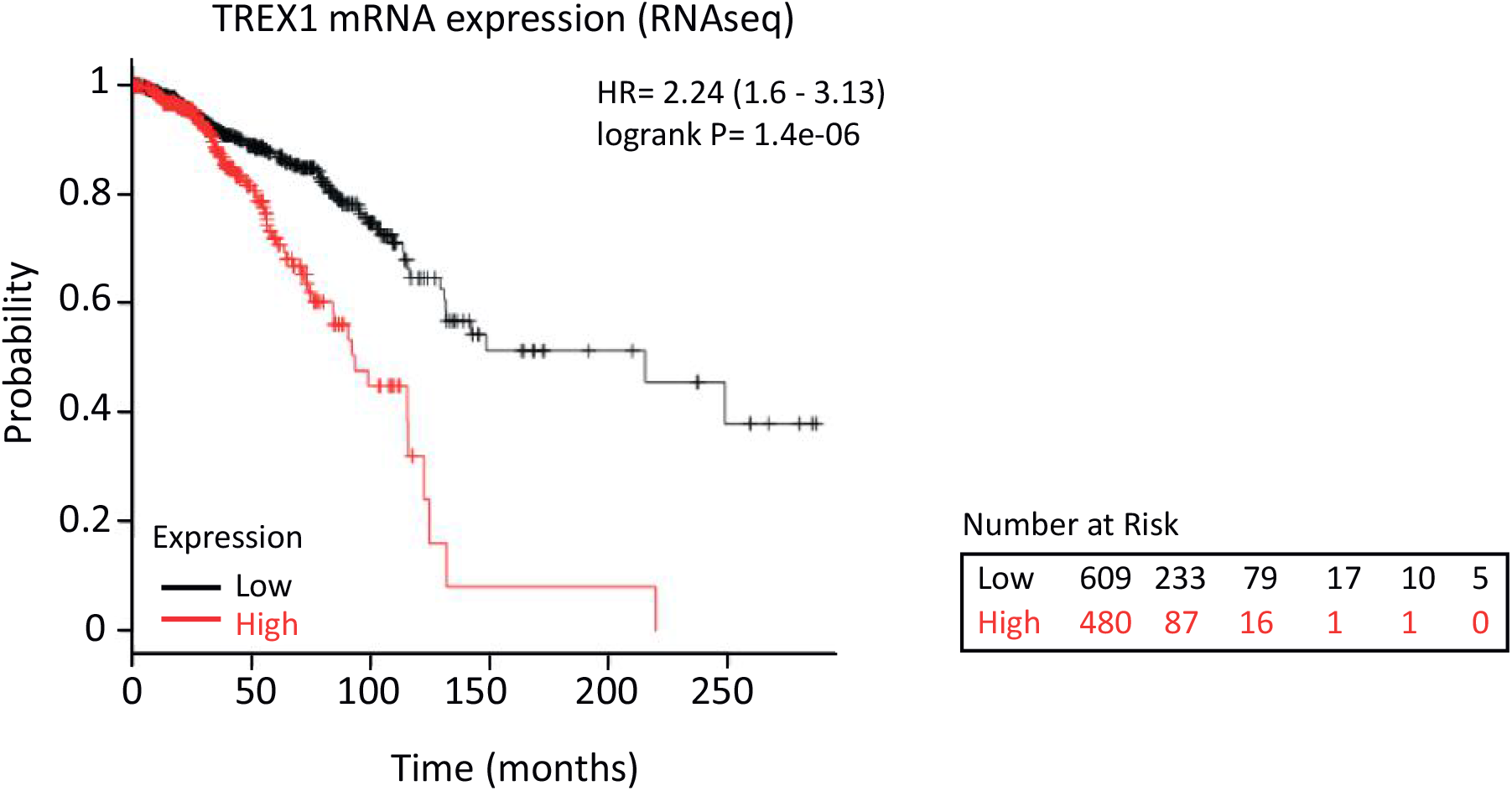
Expression of TREX1 in breast cancer is associated with decreased overall survival. Kaplan-Meier (KM) survival plot for patients expressing low (black line) and high (red line) levels of TREX1 mRNA obtained by RNAseq. Data was obtained from KM Plotter (^77^) with n=1090 TCGA breast cancer datasets. https://kmplot.com.

## Supplementary Movie Legends

**Video S1. DCIS cells moving inside the duct on chip device.** DCIS cells stably expressing 53BP1-mCherry were injected in the duct on chip and imaged under 20x magnification using spinning disc microscopy.

**Video S2. RPE1 cells exhibit repeated nuclear blebbing and rupture events under confinement.** RPE1 cells stably expressing 53BP1-EGFP and catalytically inactive cGAS-mCherry were confined at 2 μm and imaged under 40x magnification using spinning disc microscopy.

**Video S3. TREX1 exhibit nuclear localization upon confinement.** RPE1 cells were transiently transfected with TREX1-EGFP and 1 μm thickness Z slices were acquired before and after 2 μm confinement. Cells were imaged under 40x magnification using spinning disc microscopy.

**Video S4. DCIS cells subjected to strong confinement exhibit exploratory behavior in 3D collagen.** DCIS cells stably expressing life-actin-EGFP were confined at 10 μm or 2 μm for 2 h. Subsequently, cells were harvested and embedded in a 3D type I collagen matrix for imaging under 40x magnification using spinning disc microscopy.

**Video S5. Compilation of mice xenografts generated with injection of DCIS parental cells.** Three examples of tissue sections immunostained with alpha-smooth muscle actin, SMA (green), DAPI (cyan) and γH2AX (red); 6-8 weeks post-intraductal injection. Images were acquired using a spinning disc microscope (20x magnification).

**Video S6. Compilation of mice xenografts generated with injection of DCIS TREX1 KO cells.** Three examples of tissue sections immunostained with alpha-smooth muscle actin, SMA (green), DAPI (cyan) and γH2AX (red); 6-8 weeks post-intraductal injection. Images were acquired using a spinning disc microscope (20x magnification).

## References

1. Irianto, J. et al. DNA Damage Follows Repair Factor Depletion and Portends Genome Variation in Cancer Cells after Pore Migration. Curr. Biol. (2017). doi:10.1016/j.cub.2016.11.049

2. Denais, C. M. et al. Nuclear envelope rupture and repair during cancer cell migration. Science (80-.). 352, 353–358 (2016).

3. Raab, M. et al. ESCRT III repairs nuclear envelope ruptures during cell migration to limit DNA damage and cell death. Science (80-.). 352, 359–362 (2016).

4. Hatch, E. M. & Hetzer, M. W. Nuclear envelope rupture is induced by actin-based nucleus confinement. J. Cell Biol. (2016). doi:10.1083/jcb.201603053

5. Hatch, E. M. Nuclear envelope rupture: little holes, big openings. Current Opinion in Cell Biology (2018). doi:10.1016/j.ceb.2018.02.001

6. Lusk, C. P. & Ader, N. R. CHMPions of repair: Emerging perspectives on sensing and repairing the nuclear envelope barrier. Curr. Opin. Cell Biol. 64, 25–33 (2020).

7. Le Berre, M., Aubertin, J. & Piel, M. Fine control of nuclear confinement identifies a threshold deformation leading to lamina rupture and induction of specific genes. Integr. Biol. 4, 1406 (2012).

8. Gentili, M. et al. The N-Terminal Domain of cGAS Determines Preferential Association with Centromeric DNA and Innate Immune Activation in the Nucleus. Cell Rep. (2019). doi:10.1016/j.celrep.2019.03.049

9. Vargas, J. D., Hatch, E. M., Anderson, D. J. & Hetzer, M. W. Transient nuclear envelope rupturing during interphase in human cancer cells. Nucleus (2012). doi:10.4161/nucl.18954

10. De vos, W. H. et al. Repetitive disruptions of the nuclear envelope invoke temporary loss of cellular compartmentalization in laminopathies. Hum. Mol. Genet. (2011). doi:10.1093/hmg/ddr344

11. Robijns, J., Houthaeve, G., Braeckmans, K. & De Vos, W. H. Loss of Nuclear Envelope Integrity in Aging and Disease. in International Review of Cell and Molecular Biology (2018). doi:10.1016/bs.ircmb.2017.07.013

12. Cho, S. et al. Mechanosensing by the Lamina Protects against Nuclear Rupture, DNA Damage, and Cell-Cycle Arrest. Dev. Cell (2019). doi:10.1016/j.devcel.2019.04.020

13. Roman, W. et al. Myofibril contraction and crosslinking drive nuclear movement to the periphery of skeletal muscle. Nat. Cell Biol. (2017). doi:10.1038/ncb3605

14. Earle, A. J. et al. Mutant lamins cause nuclear envelope rupture and DNA damage in skeletal muscle cells. Nat. Mater. (2020). doi:10.1038/s41563-019-0563-5

15. Feng, C. et al. Cyclic mechanical tension reinforces DNA damage and activates the p53-p21-Rb pathway to induce premature senescence of nucleus pulposus cells. Int. J. Mol. Med. (2018). doi:10.3892/ijmm.2018.3522

16. Palamidessi, A. et al. Unjamming overcomes kinetic and proliferation arrest in terminally differentiated cells and promotes collective motility of carcinoma. Nat. Mater. (2019). doi:10.1038/s41563-019-0425-1

17. Lodillinsky, C. et al. p63/MT1-MMP axis is required for in situ to invasive transition in basal-like breast cancer. Oncogene (2016). doi:10.1038/onc.2015.87

18. Feinberg, T. Y. et al. Divergent Matrix-Remodeling Strategies Distinguish Developmental from Neoplastic Mammary Epithelial Cell Invasion Programs. Dev. Cell (2018). doi:10.1016/j.devcel.2018.08.025

19. Malinverno, C. et al. Endocytic reawakening of motility in jammed epithelia. Nat. Mater. (2017). doi:10.1038/nmat4848

20. Kim, M. et al. Microinvasive carcinoma versus ductal carcinoma in situ: A comparison of clinicopathological features and clinical outcomes. J. Breast Cancer (2018). doi:10.4048/jbc.2018.21.2.197

21. Atia, L. et al. Geometric constraints during epithelial jamming. Nat. Phys. (2018). doi:10.1038/s41567-018-0089-9

22. Vedula, S. R. K. et al. Emerging modes of collective cell migration induced by geometrical constraints. Proc. Natl. Acad. Sci. U. S. A. (2012). doi:10.1073/pnas.1119313109

23. Friedl, P. & Gilmour, D. Collective cell migration in morphogenesis, regeneration and cancer. Nature Reviews Molecular Cell Biology (2009). doi:10.1038/nrm2720

24. Behbod, F. et al. An intraductal human-in-mouse transplantation model mimics the subtypes of ductal carcinoma in situ. Breast Cancer Res. (2009). doi:10.1186/bcr2358

25. Blaha, L., Zhang, C., Cabodi, M. & Wong, J. Y. A microfluidic platform for modeling metastatic cancer cell matrix invasion. Biofabrication (2017). doi:10.1088/1758-5090/aa869d

26. Le Berre, M., Zlotek-Zlotkiewicz, E., Bonazzi, D., Lautenschlaeger, F. & Piel, M. Methods for Two-Dimensional Cell Confinement. in Methods in Cell Biology 213–229 (2014). doi:10.1016/B978-0-12-800281-0.00014-2

27. Shah, P., Wolf, K. & Lammerding, J. Bursting the Bubble –Nuclear Envelope Rupture as a Path to Genomic Instability? Trends in Cell Biology (2017). doi:10.1016/j.tcb.2017.02.008

28. Maciejowski, J., Li, Y., Bosco, N., Campbell, P. J. & De Lange, T. Chromothripsis and Kataegis Induced by Telomere Crisis. Cell (2015). doi:10.1016/j.cell.2015.11.054

29. Maciejowski, J., Chatzipli, A., Dananberg, A., Lange, T. de & Campbell, P. APOBEC3B-dependent kataegis and TREX1-driven chromothripsis in telomere crisis. bioRxiv (2019). doi:10.1101/725366

30. Lehtinen, D. A., Harvey, S., Mulcahy, M. J., Hollis, T. & Perrino, F. W. The TREX1 double-stranded DNA degradation activity is defective in dominant mutations associated with autoimmune disease. J. Biol. Chem. (2008). doi:10.1074/jbc.M806155200

31. Gorgoulis, V. et al. Cellular Senescence: Defining a Path Forward. Cell (2019). doi:10.1016/j.cell.2019.10.005

32. Basit, A. et al. The cGAS/STING/TBK1/IRF3 innate immunity pathway maintains chromosomal stability through regulation of p21 levels. Exp. Mol. Med. (2020). doi:10.1038/s12276-020-0416-y

33. Infante, E. et al. LINC complex-Lis1 interplay controls MT1-MMP matrix digest-on-demand response for confined tumor cell migration. Nat. Commun. (2018). doi:10.1038/s41467-018-04865-7

34. Castagnino, A. et al. Coronin 1C promotes triple-negative breast cancer invasiveness through regulation of MT1-MMP traffic and invadopodia function. Oncogene (2018). doi:10.1038/s41388-018-0422-x

35. Harding, S. M. et al. Mitotic progression following DNA damage enables pattern recognition within micronuclei. Nature (2017). doi:10.1038/nature23470

36. Lee-Kirsch, M. A. et al. A mutation in TREX1 that impairs susceptibility to granzyme A-mediated cell death underlies familial chilblain lupus. J. Mol. Med. (2007). doi:10.1007/s00109-007-0199-9

37. Lee-Kirsch, M. A. et al. Mutations in the gene encoding the 3′-5′ DNA exonuclease TREX1 are associated with systemic lupus erythematosus. Nat. Genet. (2007). doi:10.1038/ng2091

38. Lindahl, T., Barnes, D. E., Yang, Y. G. & Robins, P. Biochemical properties of mammalian TREX1 and its association with DNA replication and inherited inflammatory disease. in Biochemical Society Transactions (2009). doi:10.1042/BST0370535

39. Wolf, C. et al. RPA and Rad51 constitute a cell intrinsic mechanism to protect the cytosol from self DNA. Nat. Commun. (2016). doi:10.1038/ncomms11752

40. Ablasser, A. et al. TREX1 Deficiency Triggers Cell-Autonomous Immunity in a cGAS-Dependent Manner. J. Immunol. 192, 5993–5997 (2014).

41. Gao, D. et al. Activation of cyclic GMP-AMP synthase by self-DNA causes autoimmune diseases. Proc. Natl. Acad. Sci. U. S. A. (2015). doi:10.1073/pnas.1516465112

42. Gray, E. E., Treuting, P. M., Woodward, J. J. & Stetson, D. B. Cutting Edge: cGAS Is Required for Lethal Autoimmune Disease in the Trex1-Deficient Mouse Model of Aicardi–Goutières Syndrome. J. Immunol. (2015). doi:10.4049/jimmunol.1500969

43. Mazur, D. J. & Perrino, F. W. Excision of 3′ termini by the Trex1 and TREX2 3′→5′ exonucleases. Characterization of the recombinant proteins. J. Biol. Chem. (2001). doi:10.1074/jbc.M100623200

44. Stetson, D. B., Ko, J. S., Heidmann, T. & Medzhitov, R. Trex1 Prevents Cell-Intrinsic Initiation of Autoimmunity. Cell (2008). doi:10.1016/j.cell.2008.06.032

45. Chowdhury, D. et al. The Exonuclease TREX1 Is in the SET Complex and Acts in Concert with NM23-H1 to Degrade DNA during Granzyme A-Mediated Cell Death. Mol. Cell (2006). doi:10.1016/j.molcel.2006.06.005

46. Martinvalet, D., Zhu, P. & Lieberman, J. Granzyme A induces caspase-independent mitochondrial damage, a required first step for apoptosis. Immunity (2005). doi:10.1016/j.immuni.2005.02.004

47. Yang, Y. G., Lindahl, T. & Barnes, D. E. Trex1 Exonuclease Degrades ssDNA to Prevent Chronic Checkpoint Activation and Autoimmune Disease. Cell (2007). doi:10.1016/j.cell.2007.10.017

48. Liu, H. et al. Nuclear cGAS suppresses DNA repair and promotes tumorigenesis. Nature (2018). doi:10.1038/s41586-018-0629-6

49. Jiang, Y. N. et al. Interleukin 6-triggered ataxia-telangiectasia mutated kinase activation facilitates epithelial-to-mesenchymal transition in lung cancer by upregulating vimentin expression. Exp. Cell Res. (2019). doi:10.1016/j.yexcr.2019.05.011

50. Peng, B., Ortega, J., Gu, L., Chang, Z. & Li, G. M. Phosphorylation of proliferating cell nuclear antigen promotes cancer progression by activating the ATM/Akt/GSK3β/Snail signaling pathway. J. Biol. Chem. (2019). doi:10.1074/jbc.RA119.007897

51. Kang, H. T. et al. Chemical screening identifies ATM as a target for alleviating senescence. Nat. Chem. Biol. (2017). doi:10.1038/nchembio.2342

52. Tse, J. M. et al. Mechanical compression drives cancer cells toward invasive phenotype. Proc. Natl. Acad. Sci. U. S. A. (2012). doi:10.1073/pnas.1118910109

53. Lomakin, A., Nader, G. & Piel, M. Forcing Entry into the Nucleus. Developmental Cell (2017). doi:10.1016/j.devcel.2017.11.015

54. Lomakin, A. J. et al. The nucleus acts as a ruler tailoring cell responses to spatial constraints. bioRxiv (2019). doi:10.1101/863514

55. Enyedi, B., Jelcic, M. & Niethammer, P. The Cell Nucleus Serves as a Mechanotransducer of Tissue Damage-Induced Inflammation. Cell 165, 1160–1170 (2016).

56. Kirby, T. J. & Lammerding, J. Emerging views of the nucleus as a cellular mechanosensor. Nature Cell Biology (2018). doi:10.1038/s41556-018-0038-y

57. Elosegui-Artola, A. et al. Force Triggers YAP Nuclear Entry by Regulating Transport across Nuclear Pores. Cell 171, 1397–1410.e14 (2017).

58. Aureille, J. et al. Nuclear envelope deformation controls cell cycle progression in response to mechanical force. EMBO Rep. 20, (2019).

59. Lee, J. Y. et al. YAP-independent mechanotransduction drives breast cancer progression. Nat. Commun. 10, 1848 (2019).

60. Elster, D. et al. TRPS1 shapes YAP/TEAD-dependent transcription in breast cancer cells. Nat. Commun. (2018). doi:10.1038/s41467-018-05370-7

61. von Eyss, B. et al. A MYC-Driven Change in Mitochondrial Dynamics Limits YAP/TAZ Function in Mammary Epithelial Cells and Breast Cancer. Cancer Cell (2015). doi:10.1016/j.ccell.2015.10.013

62. Real, S. A. S. et al. Aberrant Promoter Methylation of YAP Gene and its Subsequent Downregulation in Indian Breast Cancer Patients. BMC Cancer (2018). doi:10.1186/s12885-018-4627-8

63. Nava, M. M. et al. Heterochromatin-Driven Nuclear Softening Protects the Genome against Mechanical Stress-Induced Damage. Cell (2020). doi:10.1016/j.cell.2020.03.052

64. Vanpouille-Box, C. et al. DNA exonuclease Trex1 regulates radiotherapy-induced tumour immunogenicity. Nat. Commun. (2017). doi:10.1038/ncomms15618

65. Erdal, E., Haider, S., Rehwinkel, J., Harris, A. L. & McHugh, P. J. A prosurvival DNA damage-induced cytoplasmic interferon response is mediated by end resection factors and is limited by Trex1. Genes Dev. (2017). doi:10.1101/gad.289769.116

66. Benci, J. L. et al. Opposing Functions of Interferon Coordinate Adaptive and Innate Immune Responses to Cancer Immune Checkpoint Blockade. Cell (2019). doi:10.1016/j.cell.2019.07.019

67. Takahashi, A. et al. Downregulation of cytoplasmic DNases is implicated in cytoplasmic DNA accumulation and SASP in senescent cells. Nat. Commun. (2018). doi:10.1038/s41467-018-03555-8

68. Harada, T. et al. Nuclear lamin stiffness is a barrier to 3D migration, but softness can limit survival. J. Cell Biol. (2014). doi:10.1083/jcb.201308029

69. Bell, E. S. & Lammerding, J. Causes and consequences of nuclear envelope alterations in tumour progression. European Journal of Cell Biology (2016). doi:10.1016/j.ejcb.2016.06.007

70. Pfeifer, C. R., Irianto, J. & Discher, D. E. Nuclear Mechanics and Cancer Cell Migration. in Advances in Experimental Medicine and Biology 117–130 (2019). doi:10.1007/978-3-030-17593-1_8

71. Tatli, M. & Medalia, O. Insight into the functional organization of nuclear lamins in health and disease. Current Opinion in Cell Biology (2018). doi:10.1016/j.ceb.2018.05.001

## Supplementary references

72. Lahaye, X. et al. The Capsids of HIV-1 and HIV-2 Determine Immune Detection of the Viral cDNA by the Innate Sensor cGAS in Dendritic Cells. Immunity (2013). doi:10.1016/j.immuni.2013.11.002

73. Lizárraga, F. et al. Diaphanous-related formins are required for invadopodia formation and invasion of breast tumor cells. Cancer Res. (2009). doi:10.1158/0008-5472.CAN-08-3709

74. Teullière, J. et al. Targeted activation of β-catenin signaling in basal mammary epithelial cells affects mammary development and leads to hyperplasia. Development (2005). doi:10.1242/dev.01583

75. Monteiro, P. et al. Endosomal WASH and exocyst complexes control exocytosis of MT1-MMP at invadopodia. J. Cell Biol. (2013). doi:10.1083/jcb.201306162

76. Liu, Y.-J. et al. Confinement and Low Adhesion Induce Fast Amoeboid Migration of Slow Mesenchymal Cells. Cell 160, 659–672 (2015).

77. Nagy, Á., Lánczky, A., Menyhárt, O. & Gyorffy, B. Validation of miRNA prognostic power in hepatocellular carcinoma using expression data of independent datasets. Sci. Rep. (2018). doi:10.1038/s41598-018-27521-y

